# Pan-neutralization of parainfluenza viruses by a hemagglutinin-neuraminidase antibody

**DOI:** 10.64898/2026.09.07.749649

**Authors:** Anna De Marco, Risako Gen, Young-Jun Park, Giuseppe Cusumano, Cecily Gibson, Siddhant Vyas, Corey Momont, Federico Tasin, Silvia Chiara Galli, Ben Merz, Isabella Giacchetto-Sasselli, Eneida Vetti, Tyson Rietz, Brian Tsu, Kevin Yim, Jorge Blanco, Renato Piantanida, Nadine Czudnochowski, Jessica L. Miller, Gyorgy Snell, Marina S. Boukhvalova, Jennifer E. Towne, Davide Corti, Matteo Samuele Pizzuto, David Veesler

## Abstract

Human parainfluenza viruses (HPIVs) can cause severe respiratory illnesses, such as croup, bronchiolitis, and pneumonia, particularly in children, the elderly and immunocompromised individuals. No vaccines or specific therapeutics are available for use in humans. Here, we report the discovery of a human monoclonal antibody designated PVA269 that broadly and potently neutralizes all four HPIV subtypes, PIV5, and Sendai virus by targeting the hemagglutinin-neuraminidase (HN) glycoprotein. We show that PVA269 inhibits neuraminidase activity and hemagglutination of erythrocytes through insertion of a long heavy chain complementary-determining region 3 in the enzyme active site. We reveal that the antibody markedly remodels its interactions to accommodate distinct viral features across HPIV subtypes, such as HPIV2 N-linked glycans. These results define the molecular basis for the unique PVA269 pan-neutralizing activity of human and animal viruses spanning two genera. PVA269 provides potent prophylactic activity against HPIV3 replication in the upper and lower airways of the clinically predictive cotton rat model, thus supporting translation of its protective efficacy to humans. These data establish PVA269 as a best-in-class monoclonal antibody and a promising clinical candidate to prevent HPIV infection, transmission, and disease in vulnerable populations.

## Introduction

Human parainfluenza viruses (HPIVs) are common respiratory pathogens that circulate seasonally in the human population^1^. HPIVs are part of the Paramyxoviridae family and comprise four subtypes subdivided into two genera: HPIV1 and HPIV3 (Respirovirus) and HPIV2 and HPIV4 (Rubulavirus), with HPIV4 further subdivided into HPIV4a and HPIV4b^2^. In healthy adults, HPIV infections typically cause mild, common cold symptoms in the upper respiratory tract. However, once the infection spreads to the lower respiratory tract (LRT), it can progress to severe illness such as croup, bronchiolitis, and pneumonia that mainly affect young children, the elderly, and immunocompromised individuals^3^. As a result, HPIVs are the second leading cause of hospitalization for respiratory illness in young children (after respiratory syncytial virus (RSV)), accounting for 2-17% of cases^4^. Furthermore, immunocompromised hematopoietic stem cell transplantation (HSCT) patients are at highest risk of severe disease due to early posttransplant vulnerability, immunosuppression, and comorbidity factors^5^. HSCT patients experience up to 30% HPIV infection incidence in the first 6 months following transplantation, with almost half of them progressing to LRT disease and a large fraction of them succumbing to infection^6,7^. Beyond acute infections, a growing body of evidence indicates that HPIVs and other respiratory pathogens are associated with increased prevalence of chronic disease later in life^8^. As there are no vaccines or specific antiviral treatments for any HPIVs, there is a need for therapeutics to protect these vulnerable populations.

HPIVs have two surface-exposed glycoproteins, fusion (F) and hemagglutinin-neuraminidase (HN), which work together to promote viral entry into host cells. HN is an oligomeric type II membrane protein that is responsible for engaging sialic acid-containing host receptors^9,10^, activation of F to trigger fusion of the viral and plasma membranes^11,12^, and cleaving sialic acid for virion release^13^. HN comprises a cytoplasmic tail, a transmembrane segment, a helical stalk and a b-propeller catalytic domain mediating sialoside attachment and cleavage. Despite considerable HN evolutionary distance, which can lead to amino acid sequence identities as low as 20% across HPIV subtypes, the enzymatic pocket displays remarkable conservation. This observation underscores its key functional role and makes it an ideal target for broad-spectrum inhibition of HPIVs. The HN oligomeric state seems to vary among virus subtypes, as HPIV3 HN has been shown to form dimers whereas PIV5 HN assembles as tetramers^10,14^. However, there are no available structures of HPIV1, HPIV2, or HPIV4, which hampers structure-based therapeutic design against these subtypes.

Monoclonal antibodies (mAbs) are effective prophylactics or therapeutics against respiratory viruses. For instance, nirsevimab was recently approved to be used in infants for the prevention of lower respiratory tract disease caused by RSV, showcasing the utility and success of these treatments^15^. Several HPIV neutralizing mAbs targeting F or HN have been shown to confer *in vivo* protection in small animal challenge studies but most of them are subtype-specific or characterized by narrow breadth, limiting their therapeutic _utility_^14,16–19^.

Here, we sought to identify a pan-subtype, HN-directed mAb by profiling the memory B-cell repertoire of human subjects with prior HPIV exposure. We discovered a lead candidate, designated PVA269, that cross-reacts with and inhibits all four HPIV subtypes along with animal PIVs. We determined structures of PVA269 bound to the HPIV3 head and the HPIV2 ectodomain in both dimeric and tetrameric states, revealing that PVA269 targets the conserved enzymatic pocket. In a cotton rat challenge model, we show that PVA269 provides prophylactic protection against HPIV3 challenge and reduces viral titers in both lung and nose, establishing this mAb as a promising candidate for clinical advancement to tackle HPIV infection and transmission.

## Results

### Identification of a pan-HPIV neutralizing antibody

We set out to identify a mAb cross-reacting with all four HPIV subtypes from memory B cells of convalescent human donors. To reach this goal, we analyzed the specificity of antibodies secreted by individual memory B cells (MBCs) obtained from the tonsils of a panel of human subjects, upon activation with interleukin-2 and R848 (PMID: 19404981). We detected antibodies reacting with at least one HN glycoprotein subtype in each donor and some subjects harbored MBCs specific for each of the four HN subtypes tested, likely due to prior exposure to HPIVs. However, we observed a very low frequency of MBCs cross-reacting with HPIV1-4 HN which were most enriched in the repertoire of one out of fifty donors analyzed. These findings reflect the marked evolutionary distance among HPIV subtypes with HN sharing ∼40% and ∼20% amino acid sequence identity within and across genera, respectively. The frequent detection of HPIV3 HN-directed MBCs in all donors reflects the dominant prevalence of this virus among HPIV infections.

We subsequently used fluorescence-assisted cell sorting to isolate class-switched MBCs from the donor identified above using HPIV2 HN as bait before culturing them on a mesenchymal cell monolayer with a cocktail of stimuli to promote secretion of monoclonal antibodies (mAbs) into culture supernatants. Secreted mAbs were screened for their ability to neutralize authentic HPIV3 *in vitro* and the immunoglobulin variable domains of MBCs secreting neutralizing mAbs were subsequently cloned. Our discovery campaign involved the interrogation of over 75,000 MBCs, among which HPIV2-3 cross-neutralizing activity was very rare. Of the 154 mAbs selected for further analysis, only few members of a clonal family composed of 51 mAbs (**Fig 1A and S1A, B**) fulfilled these criteria, with PVA269 being the sole member cross-reacting with and potently neutralizing all HPIV subtypes (**Fig 1B-C**). Evaluation of neutralizing activity for the 51 members of the PVA269 clonal family and the inferred germline precursor suggests that PVA269 was most likely elicited by a HPIV3 infection (**Fig 1B**). Subsequent accumulation of somatic hypermutations appears to have enhanced its potency towards the related HPIV1 subtype and enabled cross-neutralization of HPIV2 (**Fig 1B**). Consistent with this hypothesis, surface plasmon resonance (SPR) analysis demonstrated that PVA269 bound with higher affinity to the Respirovirus HPIV1 and HPIV3 HNs relative to those of Rubulavirus HPIV2 and HPIV4 (**Fig 1C and Table S1**). PVA269 is set apart from the previously described, HN-directed PIV3-23 mAb due to its unique ability to cross-react with HPIV2 HN and its greater binding affinity for all HPIV HN glycoproteins evaluated with the exception of HPIV4b for which the affinities are similar (PMID: 38858594) (**Fig. 1C and Fig S2A**). Next, we evaluated PVA269-mediated neutralization of a panel of HPIV1-4 clinical isolates and animal paramyxoviruses (**Fig. 1D and Fig S2B, C**). PVA269 efficiently neutralized all HPIV1-4 virus strains tested, and potently inhibited PIV5 and Sendai virus. In contrast, PIV3-23 did not have detectable neutralizing activity of HPIV2 isolates and exhibited reduced potency against PIV5 relative to PVA269. Collectively, these results establish PVA269 as a unique mAb with pan-PIV neutralizing activity covering human and animal isolates.

**Figure 1.**
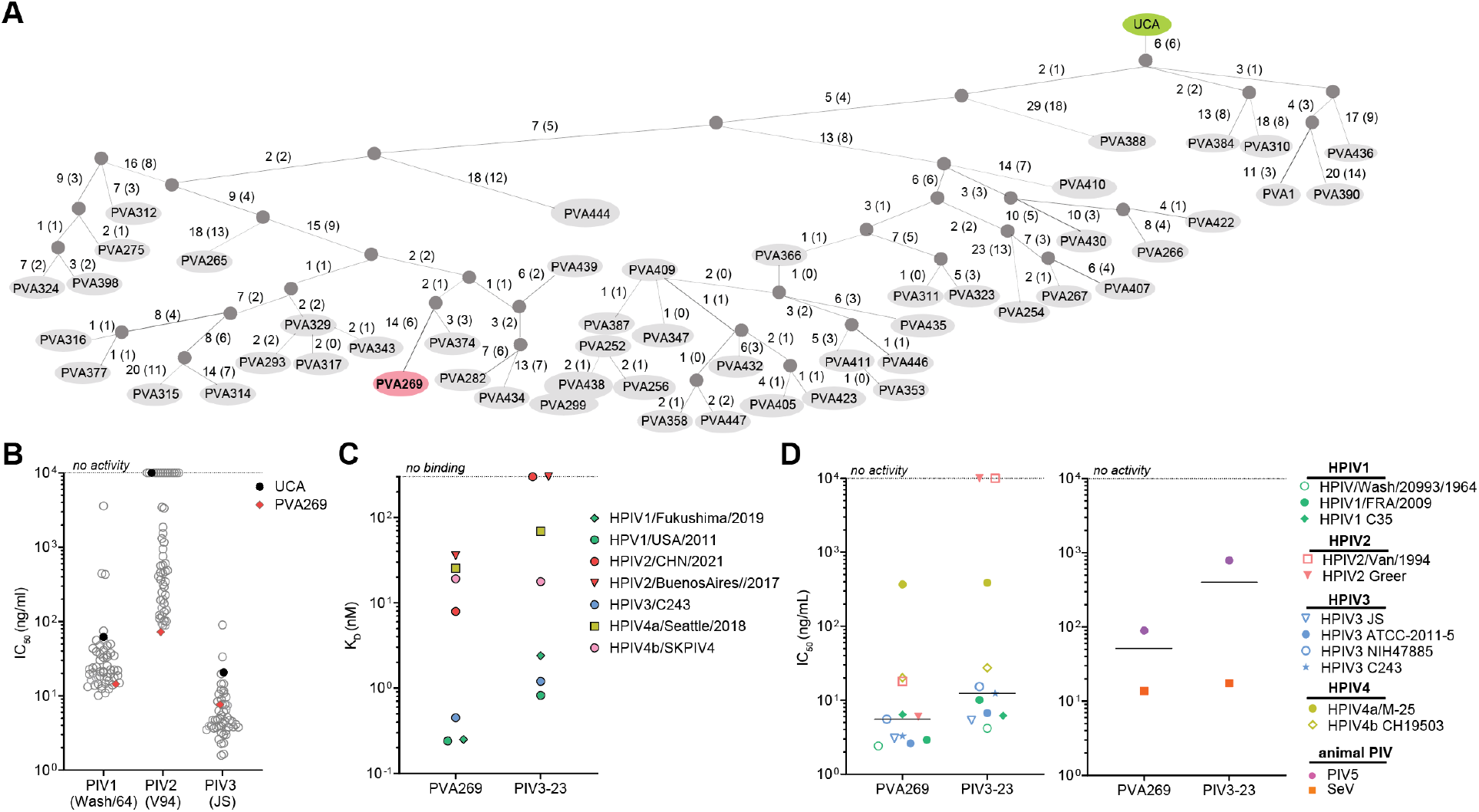
Identification of a human monoclonal antibody with potent and pan-subtype neutralizing activity. **A**. Graphical representation of the PVA VH gene clonal evolution adapted from the graphic user interface (GUI) AncesTree^20^. Branch lengths are denoted on the lines connecting the branching points with both the number of nucleotide and amino acid changes, the latter in parentheses. Branch points are indicated by grey circles. **B**. Neutralizing activity mediated by the members of the PVA269 clonal family, including the inferred germline precursor (unmutated common ancestor - UCA), against HPIV 1-3 isolates. **C**. Surface plasmon resonance binding analysis of the PVA269 and PIV3-23 mAbs to a panel of head domains (HPIV1 and HPIV3) or tetrameric HN ectodomains (HPIV2 and HPIV4). **D**. Neutralizing activity mediated by PVA269 or PIV3-23 against a panel of human HPIV 1-4 clinical isolates (left panel) and animal viruses (right panel). Conditions for which no neutralizing activity or binding were detected are shown at the limit of detection (dotted lines) in panels B-D. One independent experiment out of at least two is shown.

**Figure S1.**
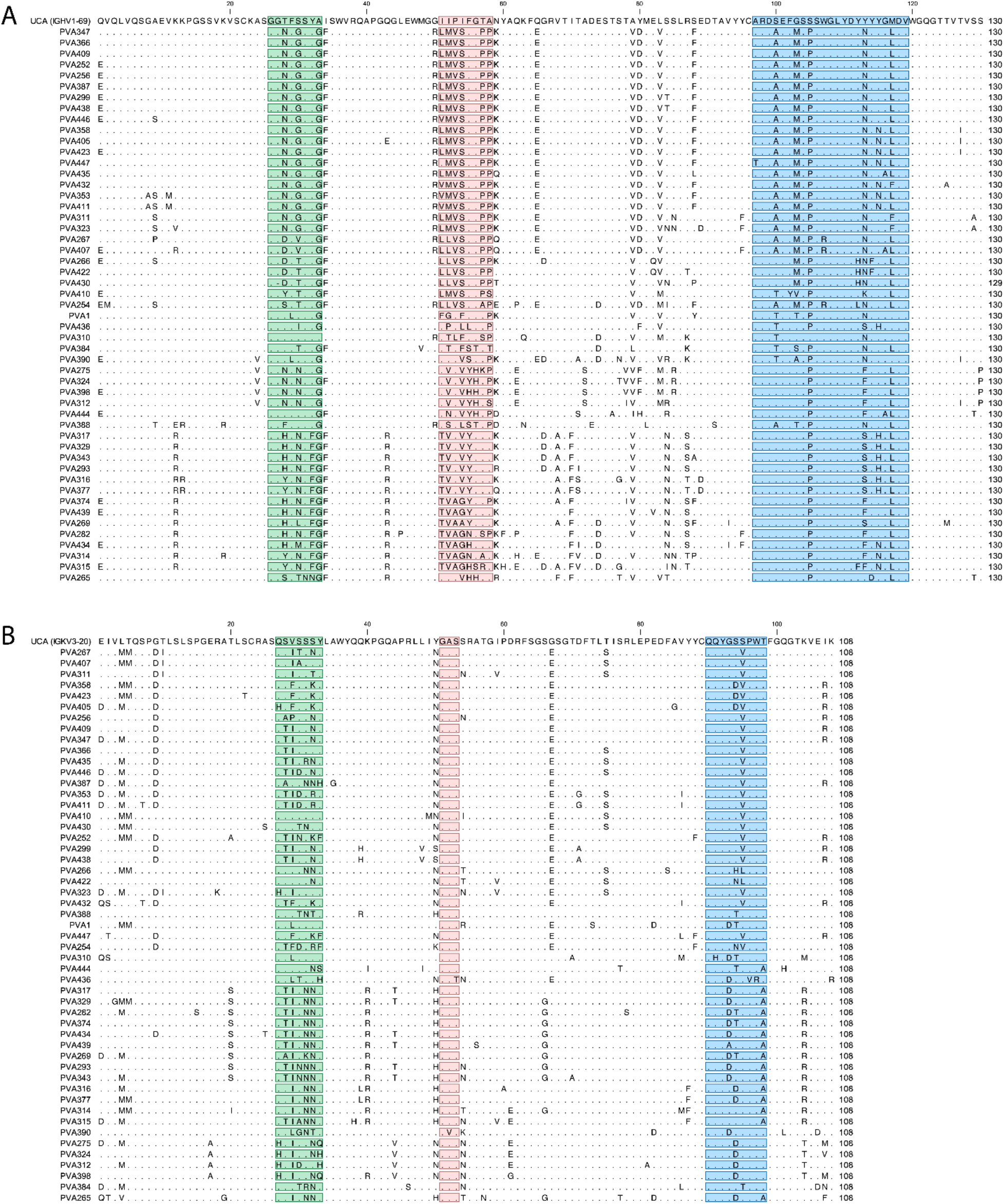
Amino acid sequence alignment of PVA269 clonal family members. Sequence alignment of the VH (A) and VL (B) genes of the 51 mAbs representing the PVA clonal family with respective unmutated common ancestors (UCAs). Complementary determining regions (CDRs) for the heavy and light chains are highlighted in green (CDR1), red (CDR2), and blue (CDR3). Residues that are identical to UCAs are shown as dots.

**Figure S2.**
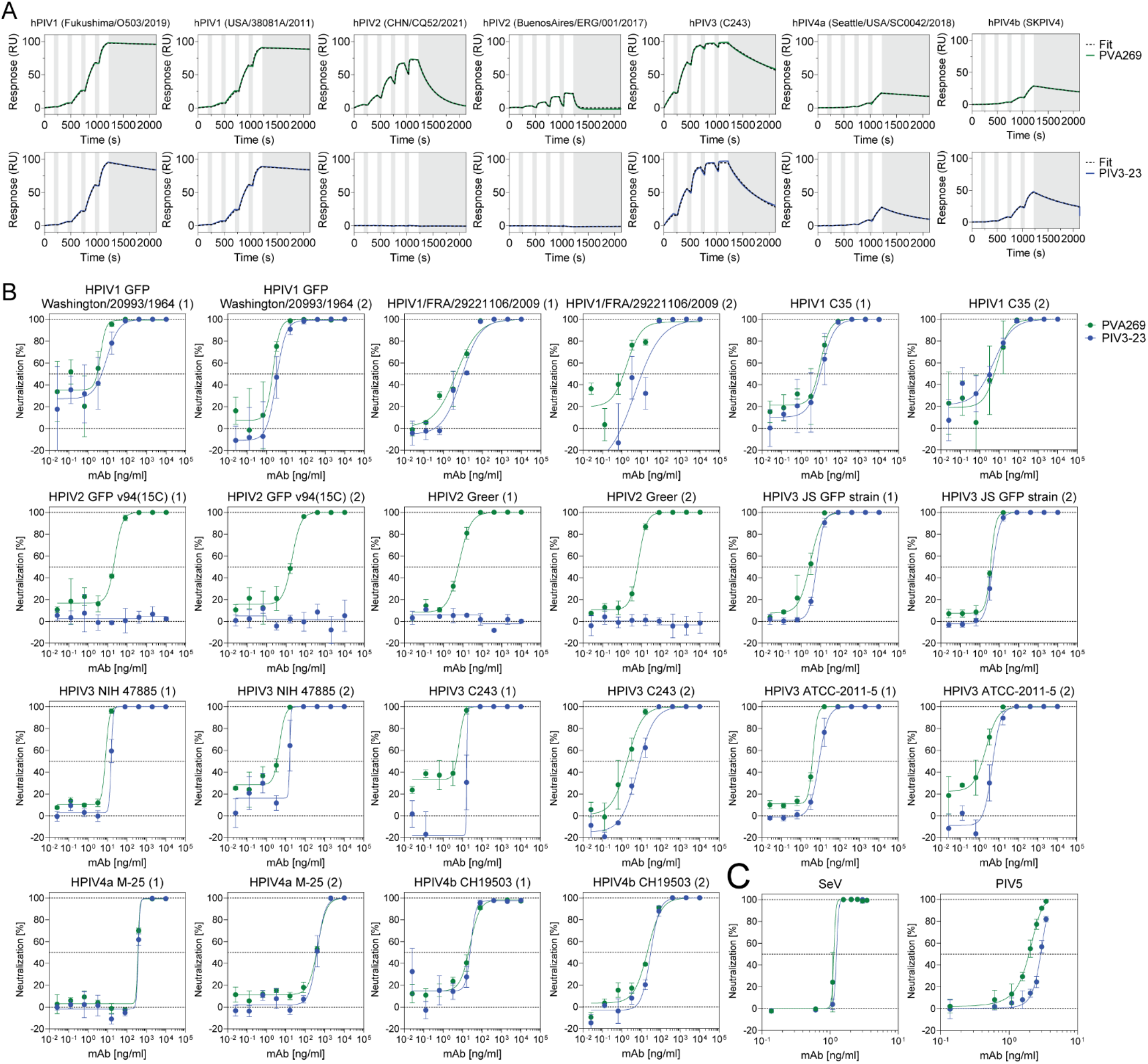
Evaluation of the PVA269 and PIV3-23 cross-reactivity and neutralization breadth. **A.** SPR sensorgrams of PVA269 or PIV3-23 recombinant Fabs binding to HPIV1-4 HN glycoproteins immobilized at the surface of SPR chips. Gray blocks denote the dissociation phases during single cycle kinetics experiments. **B,C**. Dose-response curves for the analysis of PVA269-and PIV3-23-mediated neutralization of a panel of HPIV 1-4 clinical isolates (B) and animal viruses (C) using LLC-MK2 target cells. n=2 independent biological experiments are reported for each clinical strain. Each independent experiment was performed with at least 3 technical replicates.

### PVA269 potently inhibits PIV by blocking receptor binding

To unveil the PVA269 mechanism of action, we characterized the complex between the PVA269 Fab fragment and the HPIV2 HN ectodomain using cryo-electron microscopy (cryoEM). 2D classification of the data revealed the presence of tetrameric and dimeric HN homo-oligomers, each bound to stoichiometric amounts of PVA269 Fabs, enabling the determination of cryoEM structures at 2.6Å and 2.4 Å resolution for the dimeric and tetrameric species, respectively (**Fig 2A-B, Fig S3** and **Table S2**). In the tetrameric HN structure, the four β-propeller head domains are approximately planar and form a dimer-of-dimers adopting C2 symmetry with an overall architecture reminiscent of that observed in a crystal structure of the PIV5 HN ectodomain^10^. The N-terminal stalk connecting to the viral membrane is not resolved in our structure, similar to prior HPIV3^9^ and PIV5^10^ work but distinct from another PIV5 structure^21^ as well as of paramyxovirus attachment glycoprotein Nipah virus (NiV) G^22^, Langya virus (LayV) G^23^, Newcastle disease virus (NDV) HN^24^, and Canine distemper virus (CDV) H^25^. SDS-PAGE analysis of HPIV2 HN reveals that it forms covalently linked dimers (**Fig S4A**), a property also shared with several other paramyxovirus attachment glycoproteins.

**Figure 2.**
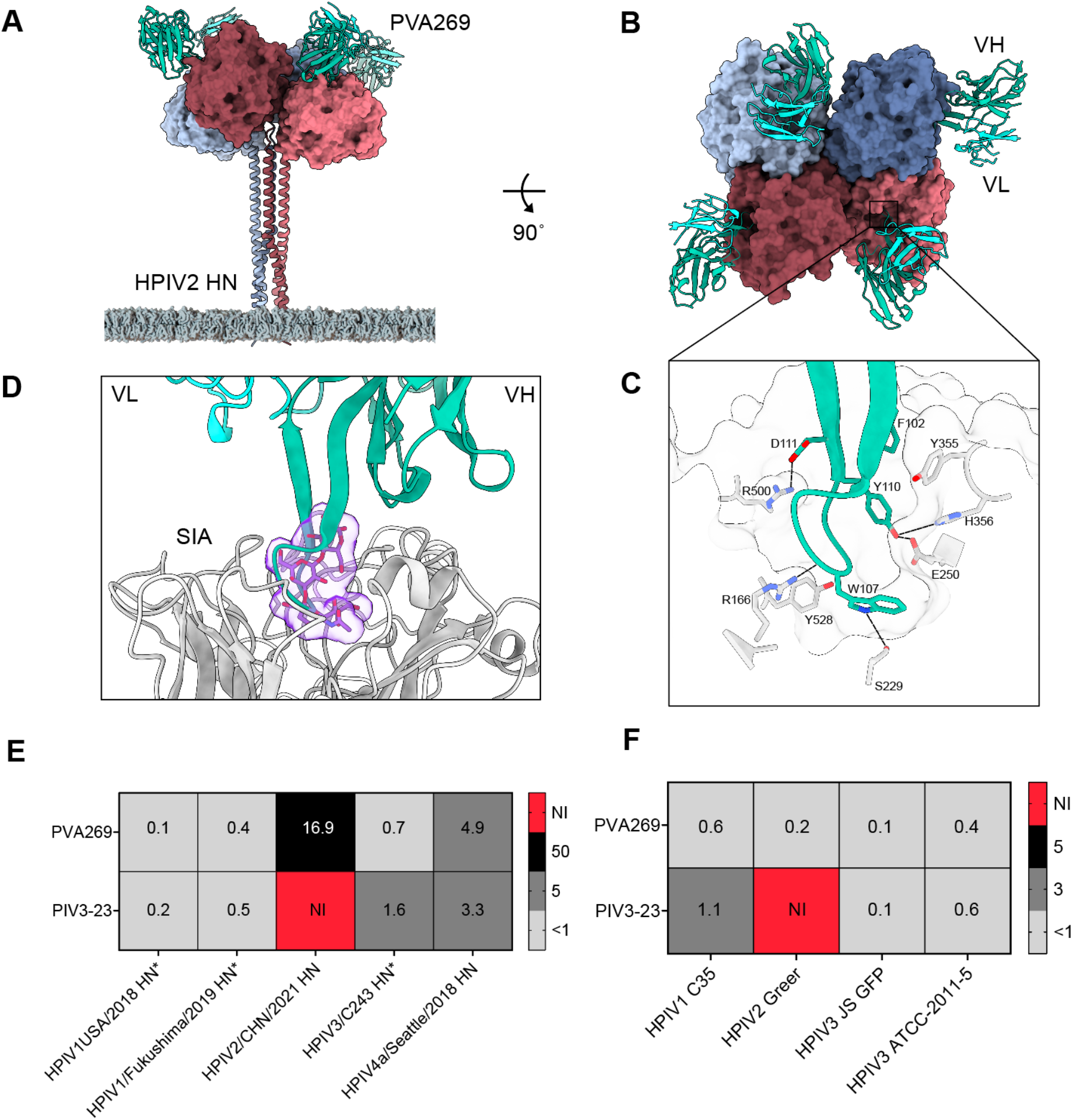
Molecular basis of PVA269-mediated pan PIV inhibition. **A-B**. CryoEM structure shown in two orthogonal orientations of the HPIV2 HN tetramer (surface representation) bound to the PVA269 Fab fragment (ribbons). The HN stalk, which is not resolved in the structure, is an AlphaFold3 predicted structure to allow visualization of the position relative to the viral membrane. Each protomer of the HN tetramer is colored distinctly and shown as a surface whereas PVA269 is rendered as dark and light green ribbons for the heavy and light chains, respectively. **C**. Zoomed-in view of the PVA269 heavy chain CDR3 projecting in the HPIV2 HN active site showing select interacting amino acid residues with polar interactions rendered as dotted lines. **D.** Superimposition of the PVA269-bound PIV2 HN structure with the sialyllactose-bound PIV5 HN structure (PDB 1Z4X) emphasizing the steric overlap between the mAb CDRH3 and the bound sialoside substrate (only the sialyllactose is shown from the PIV5 structure for clarity). **E**. PVA269-mediated inhibition (expressed as IC_50_ values in mg/ml according to the color key) of HPIV1-4 HN sialidase activity as measured using a MuNANA assay. PIV3-23 was included as a control. The reactions were carried out using the HPIV1-4 HN concentrations previously determined to generate 2×10^6^ RFU signal. NI: no inhibition detected. **F**. PVA269-mediated inhibition (IC_100_, mg/ml) of guinea pig erythrocyte hemagglutination by HPIV1-3. PIV3-23 was included as a control.

The structure reveals that the unusual, 22 residue-long heavy chain complementary determining region 3 (CDRH3) projects into the HN active site and forms a constellation of interactions, burying 660 Å^2^ at the interface with the enzyme (**Fig 2C**). Besides CDRH3 D111_PVA269_ that forms a salt bridge with R500HN, several hydrogen bonds are formed between PVA269 and HN, including between Y110_PVA269_ and E250/H356_HN_, W107_PVA269_ and R166/S229_HN_, as well as F102PVA269 and Y355_HN_, which also form T-shaped pi-stacking interactions. W107_PVA269_ and Y110 _PVA269_ make prominent contributions to the interface, respectively burying 215Å^2^ and 130Å^2^ of the paratope surface, with W107_PVA269_ positioned such that it would overlap with a bound sialoside substrate, including contacts with the conserved Y528_HN_ catalytic residue (**Fig 2C-D**). Accordingly, PVA269 inhibits the sialidase activity of purified recombinant HN glycoproteins from all four HPIV subtypes in a concentration-dependent manner, with stronger inhibition of HPIV1 and HPIV3 HNs than HPIV2 and HPIV4 HNs **(Fig. 2E and Fig. S4)**, concurring with SPR binding data (**Fig.1C**). Furthermore, PVA269 effectively blocked hemagglutination of guinea pig erythrocytes mediated by authentic HPIV1–3 virions (**Fig. 2F and Fig. S4**). Collectively, our data show that PVA269 engages the HN catalytic site and directly competes with receptor binding thus inhibiting hemagglutination of erythrocytes, neuraminidase activity and viral infectivity

**Figure S3.**
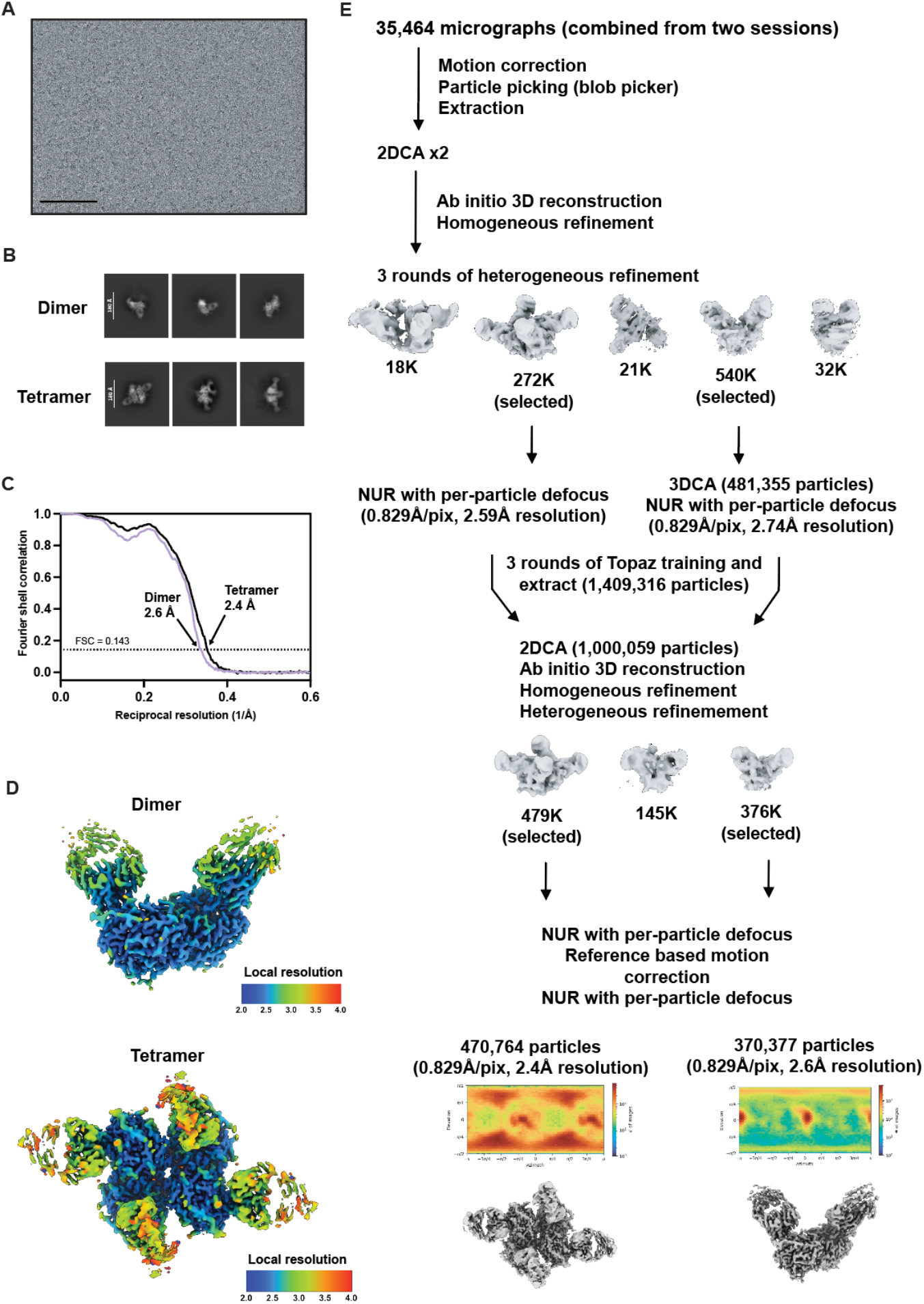
CryoEM data collection and processing of the PVA269-bound HPIV2 HN dataset. **A-B**. Representative electron micrograph and 2D class averages of the PVA269-bound HPIV2 HN dimer and tetramers embedded in vitreous ice. Scale bars: 100 nm (A) and 180 Å (B). (C) Gold-standard Fourier shell correlation curve of PVA269-bound HPIV2 dimer and tetramer structures. The 0.143 cutoff is indicated by a horizontal dashed line. (D) Local resolution estimation of the PVA269-bound HPIV2 dimer and tetramer reconstructions calculated using cryoSPARC^26^ and plotted on the corresponding sharpened maps. (E) Data processing flowchart. NUR: non-uniform refinement. The angular distribution of particle images calculated using cryoSPARC is shown as a heat map for each reconstruction.

**Fig S4.**
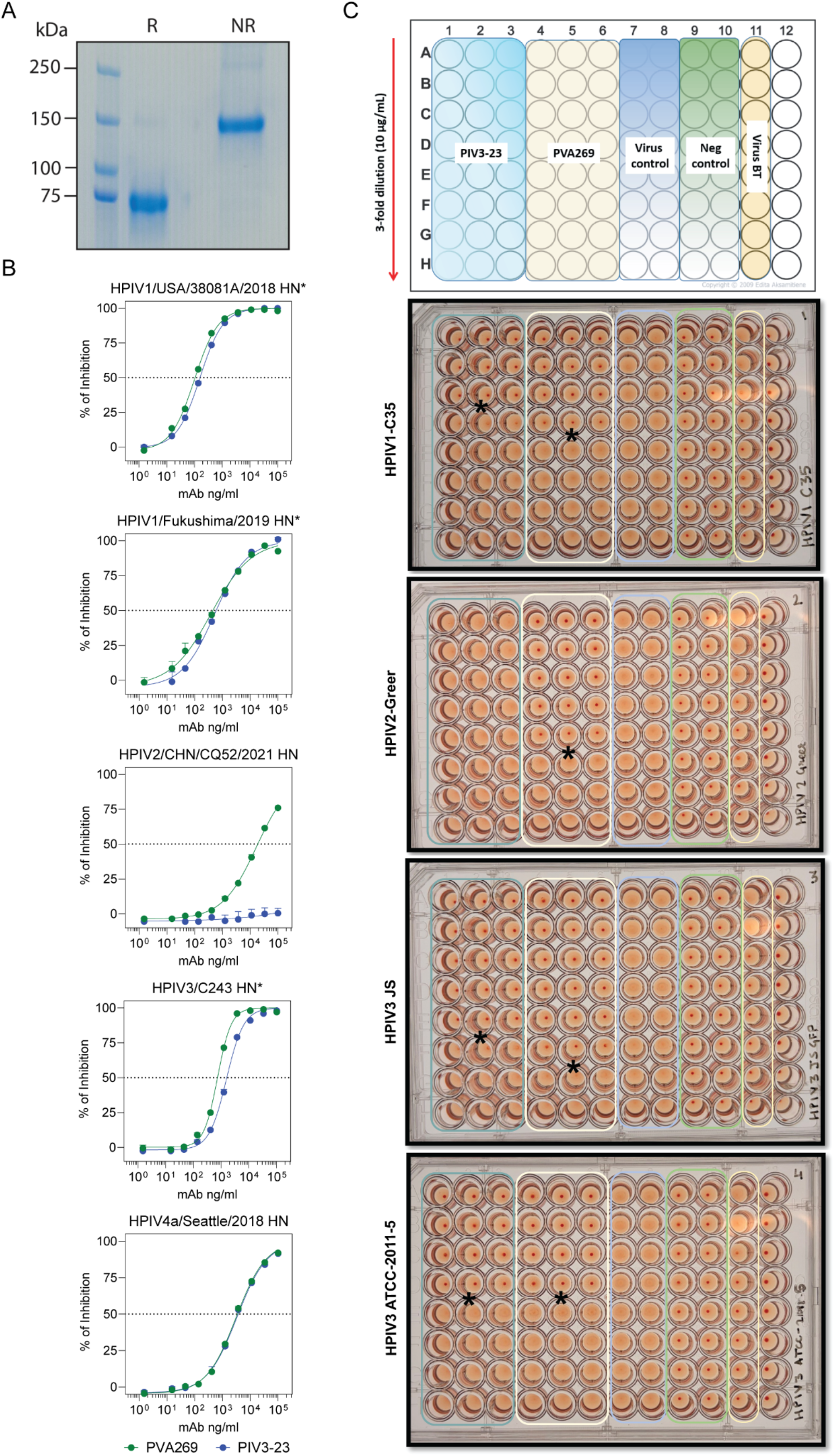
Evaluation of HN oligomerization and mAb-mediated HN inhibition. **A**. Reducing and non-reducing SDS-PAGE of the purified recombinant HPIV2 HN ectodomain used for structural studies. **B**. PVA269-and PIV3-23-mediated inhibition of HPIV1-4 HN (*head or full ectodomain) sialidase activity as measured using a MUNANA assay. **C**. PVA269-and PIV3-23-mediated inhibition of HPIV1-3 hemagglutination of guinea pig red blood cells. Asterisks mark the greatest mAb dilution at which HAI activity was observed.

### Molecular basis of PVA269-mediated pan-PIV inhibition

To understand the unique PVA269 ability to inhibit all human and animal PIV isolates evaluated, we determined a crystal structure of the HPIV3 head domain bound to the PVA269 Fab fragment at 1.68Å resolution (**Fig 3A** and **Table S3**). Comparison of the PVA269-bound and of a previously described PIV3-23-bound HPIV3 HN structures reveals conserved interactions with the active site with eight out of nine buried CDRH3 residues being identical between the two mAbs (**Fig. S5A**). Accordingly, PIV3-23 inhibited the sialidase activity of HPIV1, HPIV3, and HPIV4 HNs and blocked HPIV1-and HPIV3-mediated hemagglutination of guinea pig erythrocytes (**Fig. 2E, F and Fig. S4**). However, (**Fig. 1C,D**), PIV3-23 neither inhibited the HPIV2 HN sialidase activity nor prevented HPIV2-mediated erythrocyte hemagglutination. These results suggest that the unique pan-HPIV neutralizing activity of PVA269 is not solely driven by engagement of the HN catalytic site.

**Figure 3.**
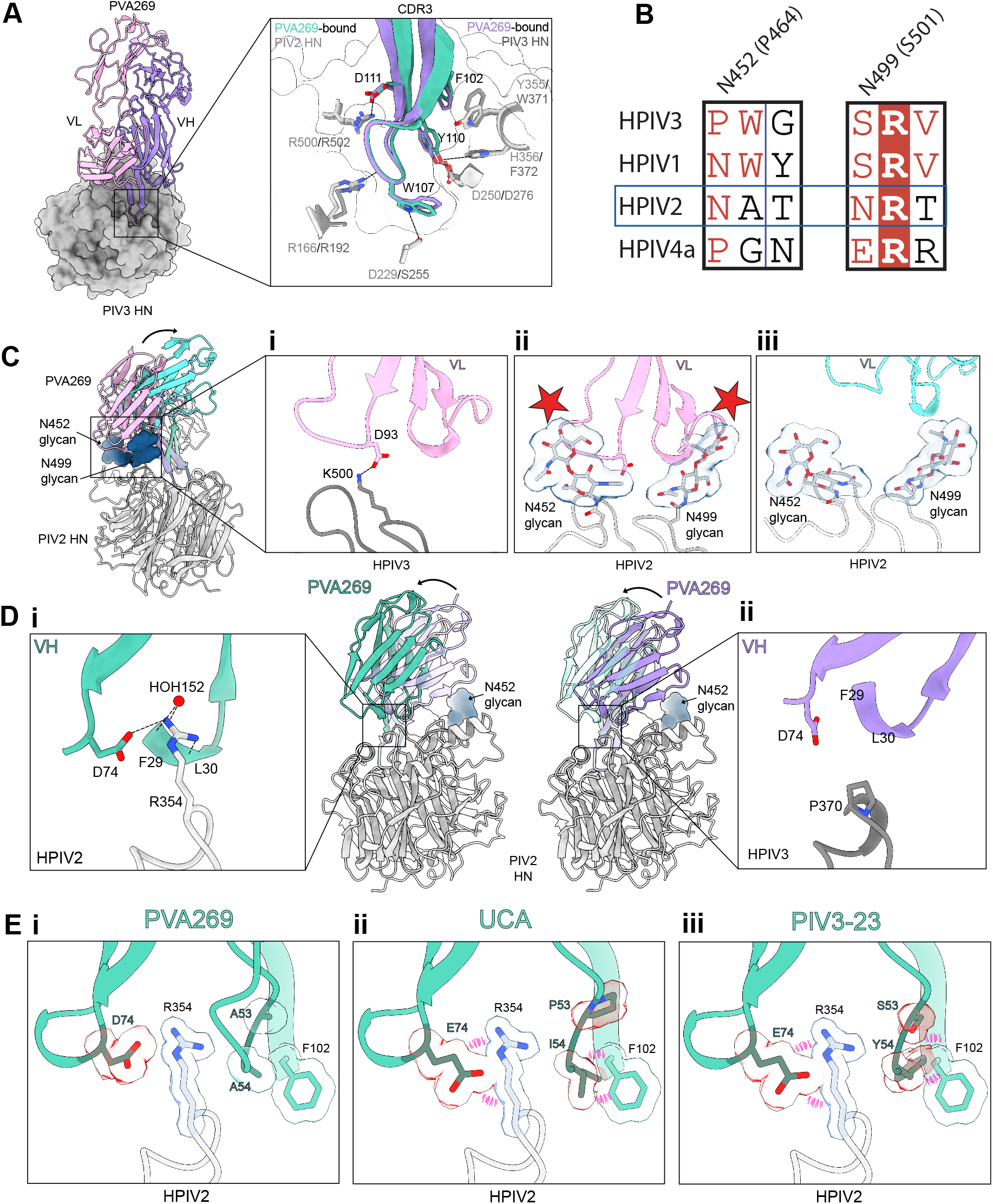
Molecular basis of PVA269-mediated pan-PIV inhibition. **A.** Crystal structure of the HPIV3 HN head (gray surface) bound to the PVA269 Fab (purple and pink for heavy and light chains, respectively). Inset: zoomed-in view of PVA269 CDRH3 from the HPIV2-(green) and HPIV3-bound (purple) structures. Select interacting residues are labeled in light grey for HPIV2 HN and dark grey for HPIV3 HN. **B.** Sequence alignment of HPIV HN glycoproteins emphasizing the presence of N-linked glycans at positions 452 and 499 that are unique to HPIV2 and are in the vicinity of the PVA269 epitope. Residue numbering shown in parentheses corresponds to HPIV3 HN. **C**. Relative positioning of the PVA269 light chain in the HPIV2-(light blue) and HPIV3-bound (pink) HN structures. Only the HPIV2 HN head (ribbons) is shown for clarity. The curved arrow indicates the relative movement between the light chains. Insets: zoomed-in views of (i) a key PVA269 light chain interaction formed with HPIV3 HN, (ii) the predicted steric clash (red stars) that would occur between PVA269 (as bound to HPIV3 HN) with HPIV2 HN glycans N452 and N499, and (iii) accommodation of HPIV2 HN glycans due to antibody repositioning as observed in the cryoEM structure. N-linked glycans near the epitope are shown as blue surfaces. **D**. Relative positioning of the PVA269 heavy chain in the HPIV2-(green) and HPIV3-bound (purple) HN structures. The curved arrows show the relative movement between the heavy chains. For clarity, the HPIV3 HN-bound PVA269 (left) or the HPIV2 HN-bound PVA269 (right) structures are shown as semi-transparent ribbons. Insets: zoomed-in views of (i) the extensive interaction network of HPIV2 HN R354 with PVA269, and (ii) lack thereof due to substitution to P370 at the equivalent HPIV3 HN position. Dash lines indicate salt bridge or hydrogen-bonding interactions. **E**. Structural analysis of PVA269 contacts formed with HPIV2 HN R354 (i) and of predicted contacts formed between the inferred, unmutated germline precursor (UCA, ii) or of PIV3-23 (iii) with HPIV2 HN R354 explaining the unique properties of PVA269. Pink dash lines indicate steric clash. Select side chains are shown in surface representation for emphasis.

Comparison of the PVA269-bound HPIV2 HN (cryoEM) and HPIV3 HN (crystal) structures show that CDRH3 residues 103-111 maintain virtually identical contacts with the HN active sites (**Fig 3A**). However, we observed a large-scale repositioning (up to 18Å shift) of the rest of the Fab relative to the head domain due to the presence of two N-linked glycans at HPIV2 HN positions N452 and N499, which would be sterically incompatible with the positioning of PVA269 observed in the HPIV3 HN-bound structure **(Fig 3B-Cii**). This results in profound remodeling of the interactions formed outside the active site. The Fab light chain buries 200Å^2^ at the interface with HPIV3 HN, including D93_PVA269_ forming a salt bridge with HPIV3 K500_HN_ (**Fig 3Ci**), whereas the HPIV2 N452_HN_ and N499_HN_ glycans prevent any interactions besides tenuous contacts with the latter oligosaccharide. On the opposite side of the epitope, HPIV2 R354_HN_ extensively interacts with the Fab heavy chain, including a salt bridge with the D74_PVA269_ side chain and two hydrogen bonds with the backbone carbonyls of F29_PVA269_ (through a water molecule) and of L30_PVA269_ (directly), burying 175Å^2^ at the interface with the paratope **(Fig 3Di**). These interactions do not occur with HPIV3 due to the inability of P370_HN_ (at the equivalent position to HPIV2 R354_HN_) to form such contacts **(Fig 3Dii**). Analysis of amino acid residue conservation of the HPIV2 and HPIV3 epitopes recognized by PVA269 indicates a strict conservation within each viral subtype, concurring with the broad cross-reactivity and neutralizing activity observed for this mAb (**Fig 1B-D, Fig S2A-B and Fig S5B**). Overall, the PVA269 heavy chain forms comparably extensive interactions with HPIV3 and HPIV2 HN, whereas its light chain interacts more strongly with HPIV3 than HPIV2 HN, likely explaining the enhanced binding affinity as well as inhibition of neuraminidase activity, hemagglutination, and *in vitro* neutralization potency observed against HPIV3 relative to HPIV2 **(Fig 1C and Fig 2Ci**).

To identify the molecular determinants setting PVA269 apart from other known antibodies in its ability to neutralize HPIV2 HN, we compared PVA269 to other sequences of the same clonal family (**Fig 1A**). Whereas CDRH3 is highly conserved to form extensive interactions with the substrate binding site, CDRH1 and CDRH2 are variable. We identified two hotspots of hypervariable Fab residues at the R354_HN_-interaction site: residues 53-54_PVA269_, and residue 74_PVA269_ (**Table S4**). Clonal family members with the most potent HPIV2 neutralizing activity harbor an aspartate residue at position 74, forming a salt bridge with R354_HN_, and small residues at positions 53-54, such as A53/G54 (PVA439) or A53/A54 (PVA269), which participate in shaping the R354_HN_-binding pocket (**Fig 3Ei**). Substitutions of these residues to a glutamate (position 74) or to bulky side chains (positions 53-54) are associated with reduced or even abrogation of HPIV2 neutralizing activity, likely due to hindering interactions with HPIV2 R354_HN_. These HPIV2 HN-specific interactions appear thus critical to compensate for the marked reduction of the PVA269 light chain contribution to the paratope. Accordingly, residues E74 and P53/I54 of the unmutated, inferred germline precursor (UCA) would remodel and disrupt the R354_HN_-binding pocket, explaining the lack of HPIV2 neutralization, which was acquired through somatic hypermutations (**Fig 1B and Fig 3Eii**). The observed retention of HPIV2 neutralizing activity (albeit lower than that of PVA269) for clonal family members harboring either the D74E mutation or substitutions introducing bulky side chains at positions 53-54 indicates that a single type of alteration can be accommodated at a time but that the combination of both changes is disruptive and abolishes interactions with R354_HN_ (**Table S4**).

As aforementioned, PVA269 and PIV3-23 share virtually identical CDRH3 and form indistinguishable contacts with the HN active site despite using distinct inferred germline genes (IGHV4-31 and IGHV1-69, respectively). However, PIV3-23 does not neutralize HPIV2 likely because it harbors E74_PIV3-23_ and a bulky tyrosine residue at position 54_PIV3-23_, which would clash with F102_PIV3-23_, thereby remodeling the key HPIV2 R354_HN_-binding pocket (**Fig 3Eiii**). To validate the role of HPIV2 HN residue R354 for interacting with PVA269, we used BLI to assess PVA269 binding to wildtype and R354A HPIV2 HN glycoprotein ectodomains. We found that the PVA269 Fab recognized immobilized wildtype HPIV2 HN with an approximate affinity of 16 nM, whereas no binding was detected to the R354A mutant up to a concentration of 160 nM. However, PVA269 IgG retained binding to both wildtype and R354A HPIV2 HN constructs, with respective approximate avidities of 16 pM and 667 pM, underscoring the crucial roles of HPIV2 R354 for PVA269 binding and of bivalent mAb binding to promote resilience to escape mutations, as previously observed in the context of trimerization of coronavirus minibinders^27–29^ (**Fig S5C-F**). Collectively, these findings provide a molecular blueprint for understanding the unique ability of PVA269 to mediate pan-PIV neutralization.

**Fig S5.**
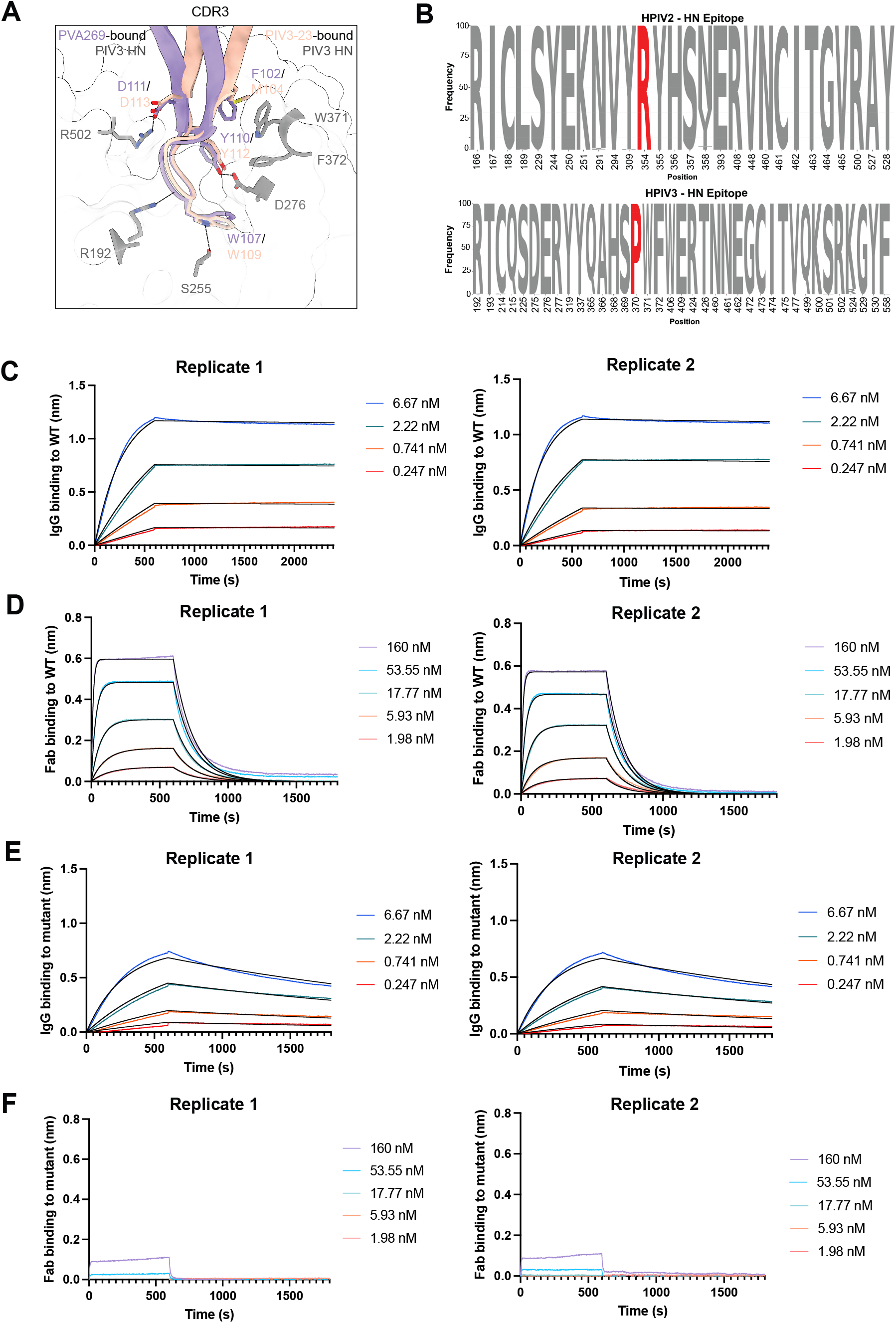
Comparison of PVA269 and PIV3-23 binding to HN, epitope conservation and evaluation of PVA269 binding to wildtype and mutant HPIV2 HN. **A.** Zoomed-in view of PVA269 (purple) and PIV3-23 (orange, PDB: 8TQI) CDRH3 from their respective HPIV3-bound structures. Interacting residues are labeled in purple for PVA269, orange for PIV3-23, and dark grey for HPIV3. **B** Logo plot amino acid conservation of PVA269 epitope based on publicly available HN sequences from HPIV2 (n = 210) and HPIV3 (n = 2140). HPIV2 residue R354 and HPIV3 residue P370 are indicated in red. **C-F**. BLI binding analysis of PVA269 IgG (C) or Fab (D) to immobilized wildtype HPIV2 HN or of PVA269 IgG (E) or Fab (F) to immobilized R354A HPIV2 HN. Two biological replicates carried out with independently produced batches of HPIV2 HN are shown. The sensorgrams are colored according to the color key and the fit to the data using a 1:1 global fitting model is shown in black.

### PVA269 protects cotton rats against HPIV3 challenge

To assess the protective efficacy of PVA269, we administered the antibody intramuscularly to cotton rats (*Sigmodon hispidus*) at doses of 3, 1.5, 0.75, 0.375, 0.1875 mg/kg one day before intranasal challenge with 10^5^ p.f.u. of HPIV3 strain C243 (ATCC VR-93, Manassas, VA) (**Fig 4A**). Quantification of HPIV3 replicating virus titers 4 days post infection showed that PVA269 reduced viral load in the lungs and in the nose in a dose-dependent manner, relative to an isotype control antibody (**Fig 4B-C**). Indeed, administration of PVA269 at 3 and 1.5 mg/kg reduced lung and nose viral titers by 1.5-2 orders of magnitudes, reaching the lower limit of detection in the nose (**Fig 4A-B**). Furthermore, an antibody dose as low as 0.75 mg/kg resulted in a marked and statistically significant reduction of replicating lung and nose viral titers in comparison to the control group, underscoring the potent PVA269 inhibitory activity. The serum mAb concentration measured one day after administration (2 hours before infection) inversely correlated with HPIV3 replication in the lungs and nose (**Fig 4D-E)**. Based on average viral titers in the control group (0% efficacy) and the assay limit of detection (LOD; 100% efficacy), the estimated PVA269 serum concentrations required to achieve 50% (EC_50_) and 99% (EC_99_) protection were 11 and 54 µg/mL in the lung and 8 and 38 µg/mL in the nose, respectively (**Fig. S6**).

**Figure 4.**
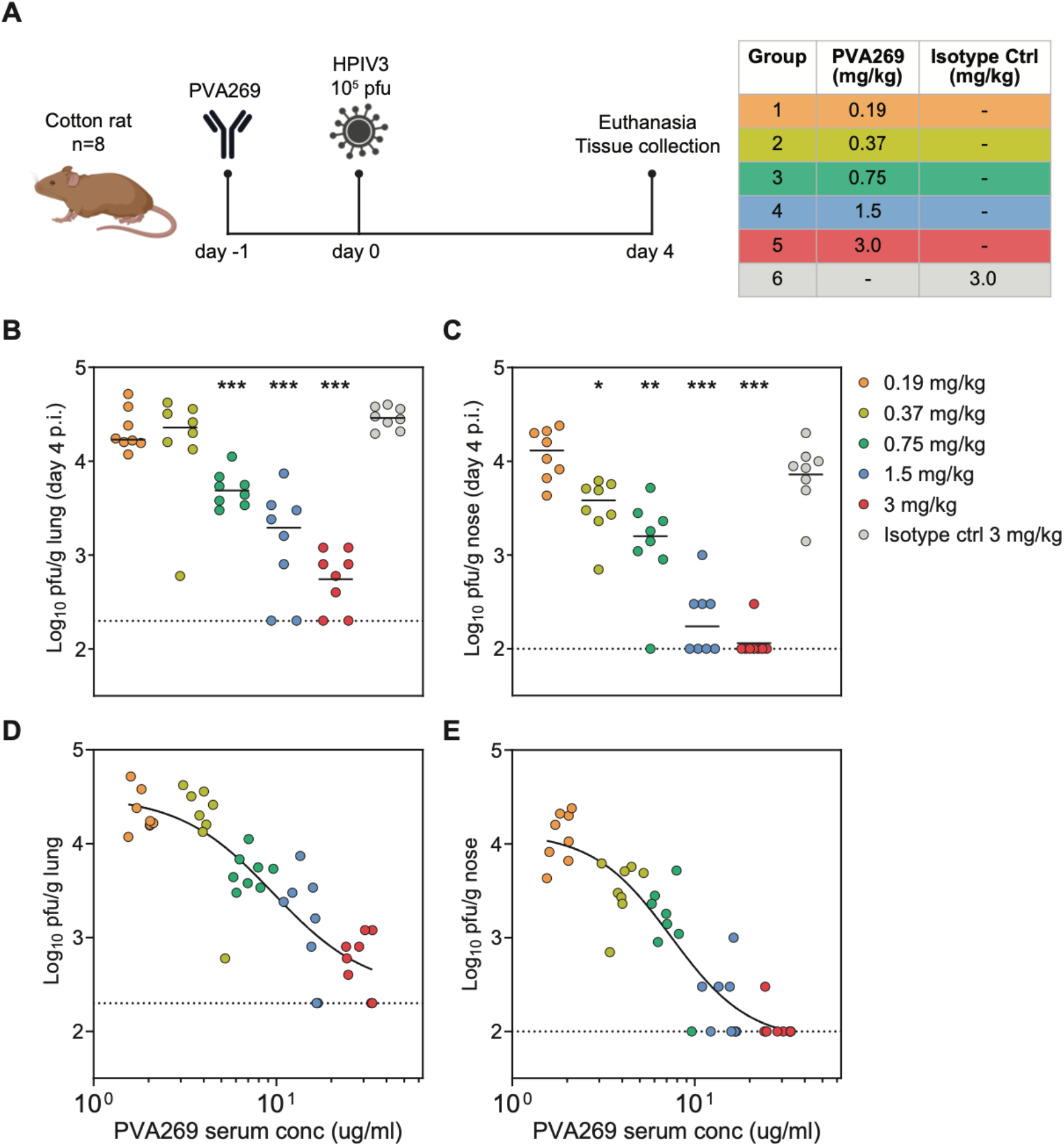
PVA269 provides prophylactic protection against HPIV3 in the upper and lower airways. **A**. Schematic of the study design for evaluation of the *in vivo* efficacy of PVA269 against HPIV3 strain C243 (ATCC VR-93, Manassas, VA) challenge in cotton rats. **B-C**. Replicating virus titres in the lungs (B) and in the nose (C) of cotton rats (n=8/group) measured 4 days post-infection. **D-E**. Efficacy curves correlating PVA269 serum titers 24 hours post-administration (i.e. on the day of challenge) with replicating viral titers in the lung (D) and in the nose (E) calculated using GraphPad Prism v10.0 using a nonlinear regression analysis.

These results indicate that PVA269 is highly efficacious in preventing replication of HPIV3 both in the upper and the lower respiratory tracts of cotton rats, establishing a direct correlation between the anti-HN mAb activity and *in vivo* protection.

**Fig S6.**
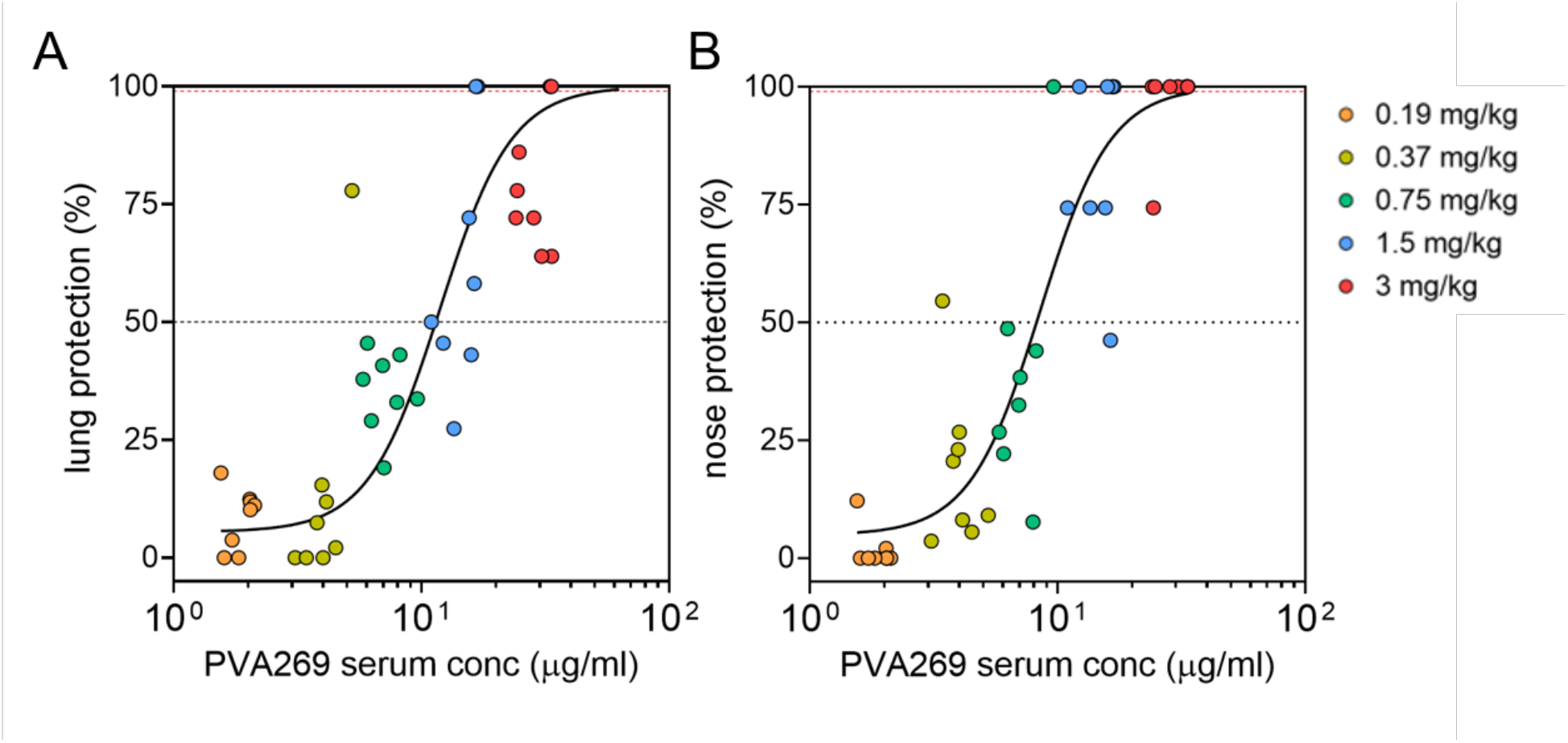
Correlation between PVA269 serum concentrations and protection of cotton rats against HPIV3 challenge. **A-B**, Non-linear regression fitting of lung (A) and nose (B) protection (derived from replicating viral titers) in function of PVA269 serum levels for the animal groups used in the challenge study. EC_50_ and EC_99_ are shown as black and red dotted lines, respectively. Replicating viral titers (protection) observed for the isotype-control antibody and the limit of detection of the plaque assays were used as 0% and 100% reference for data normalization, respectively.

## Discussion

Management of current HPIV infections is largely limited to supportive care, as no virus-specific antiviral therapies or vaccines have been approved. Aerosolized or systemic ribavirin administration has been used off-label in some severely immunocompromised patients, to limit progression to lower respiratory tract disease, but its efficacy remains uncertain. DAS181, a recombinant sialidase fusion protein designed to remove sialic-acid receptors required for viral entry, is currently being evaluated in a Phase 3 clinical trial for the treatment of lower respiratory tract HPIV infection in immunocompromised patients (NCT03808922). Moreover, vaccine development efforts have focused largely on HPIV3, given its major contribution to severe bronchiolitis and pneumonia in young infants, immunocompromised individuals, and transplant recipients, thereby limiting the expected breadth of immunity elicited. Following the clinical success of anti-RSV therapeutics, several preclinical studies described F-directed HPIV3 mAbs and showed that they reduce viral burden in small animal challenge models^14,18,30,31^. However, these mAbs are endowed with neutralizing activities of narrow breadth, underscoring the need for broad spectrum HPIV countermeasures.

Paramyxovirus attachment glycoproteins have been observed in two distinct oligomeric states. While tetramers of PIV5 HN^21^, NDV HN^24^, CDV H^25^, NiV G^22^, and LayV G^23^ have been previously described, HPIV3 HN^32^ and Measles virus (MeV) H^33^ have been observed as dimeric structures. The present identification of HPIV2 HN dimers and tetramers within the same dataset, recapitulating the previously observed architectures for other paramyxoviruses, unifies these prior observations. An average area of approximately 1,700 Å² is buried at the interface between HPIV2 HN protomers in the dimer. In contrast, an average surface smaller than 300 Å² is involved in each of the two interfaces that drive HPIV2 HN tetramerization through a dimer-of-dimers configuration. Furthermore, the cryo-EM density points to potential interactions among residues directly N-terminal to the head domains; these residues likely form a C114 interprotomer disulfide bond linking the exact same two protomers that constitute the primary dimer, concurring with the detection of a prominent dimeric species via non-reducing SDS-PAGE. This topology is reminiscent of the PIV5 HN C111 disulfide bond linking two protomers with the same head organization as that of the HPIV2 HN dimer. Overall, these results suggest that these distinct conformations and oligomeric species may represent functionally relevant oligomers for multiple HPIV subtypes and paramyxoviruses in general.

The PVA269 mAb described here was isolated from tonsillar B cells of a human donor characterized by an immune repertoire enriched with cross-reactive monoclonal antibodies, a particularly rare feature amongst the cohort of individuals investigated. Furthermore, PVA269 is a part of a larger clonal family, composed of 51 members, among which only few were able to cross-react and neutralize all four HPIV subtypes, with PVA269 standing out as the best candidate based on potency and breadth. Originating from an inferred germline precursor with narrow cross-reactivity, PVA269 achieves pan-parainfluenzavirus neutralizing activity through the accumulation of somatic hypermutations, which contributed to the breadth expansion and potency improvement, extending beyond human viruses to other animal parainfluenza viruses. PVA269-mediated viral inhibition is exerted via insertion of the long antibody HCDR3 in the HN active site, outcompeting sialoside receptors. Furthermore, PVA269 remodels its interactions with the HPIV2 HN head domain outside the active site, explaining its unique ability to retain strong binding to and neutralizing activity against viruses of the HPIV2 subtype, setting it apart from any other known HPIV mAbs. We note that PVA269 is reminiscent of the neuraminidase-directed FNI9^34^ and 1G01^35^ mAbs, which also target the enzyme active site to achieve broad neutralizing activity and protection against a wide spectrum of influenza viruses.

Cotton rat (*Sigmodon hispidus*) is considered the “gold standard model” to assess RSV-targeting and was previously used to evaluate the activity of HPIV-directed mAbs^14,31^. This animal model is particularly informative as it supports viral replication in both the upper and lower respiratory tract without prior adaptation and recapitulates key aspects of human disease, including quantifiable viral burden and pulmonary pathology. For instance, a protective serum antibody threshold was determined based on protection (EC_99_) against lung RSV loads in cotton rats to inform clinical dose selection for the palivizumab prophylactic mAb. Furthermore, cotton rats were used to benchmark the increased *in vivo* potency of nirsevimab, relative to palivizumab, in support of the once-per-season dosing strategy of the former mAb^36,37^. Using a similar experimental design, Boonyaratanakornkit *et al.* evaluated the *in vivo* efficacy of the HPIV-3 F-targeting mAb PI3-E12, which was found to prevent viral replication in the lungs with an EC_50_ of 0.35 mg/kg and an EC99 of 1.80 mg/kg^31^. Conversely, administration of PI3-E12 at 1.25 mg/kg and higher doses modestly reduced HPIV3 replication in the nose, and protection plateaued at approximately one order of magnitude reduction relative to the control group^31^. When tested in an identical setting, PVA269 provided potent protection against viral replication in the lung, with EC_50_ and EC_99_ of 11 and 54 µg/ml of mAb in the serum, respectively. Furthermore, viral replication in the upper respiratory tract (nose) reached the lower limit of detection in half of the animals (4/8) when PVA269 was administered at 1.5 mg/kg and in all but one animal (7/8) in those receiving the antibody at 3 mg/kg. These results demonstrate that the pan-parainfluenza, HN-directed PVA269 mAb confers potent protection in the upper and lower respiratory tract in a clinically predictive model, supporting its potential to protect humans efficiently. We anticipate that combining the breadth and potency of PVA269 with Fc-mediated half-life extension could enable season-long protection, reducing the burden of HPIV disease and potentially limiting transmission. Collectively, our findings establish PVA269 mAb as a best-in-class candidate with the potential to transform prophylactic strategies against HPIV infection in vulnerable populations.

---

**Table S1.**
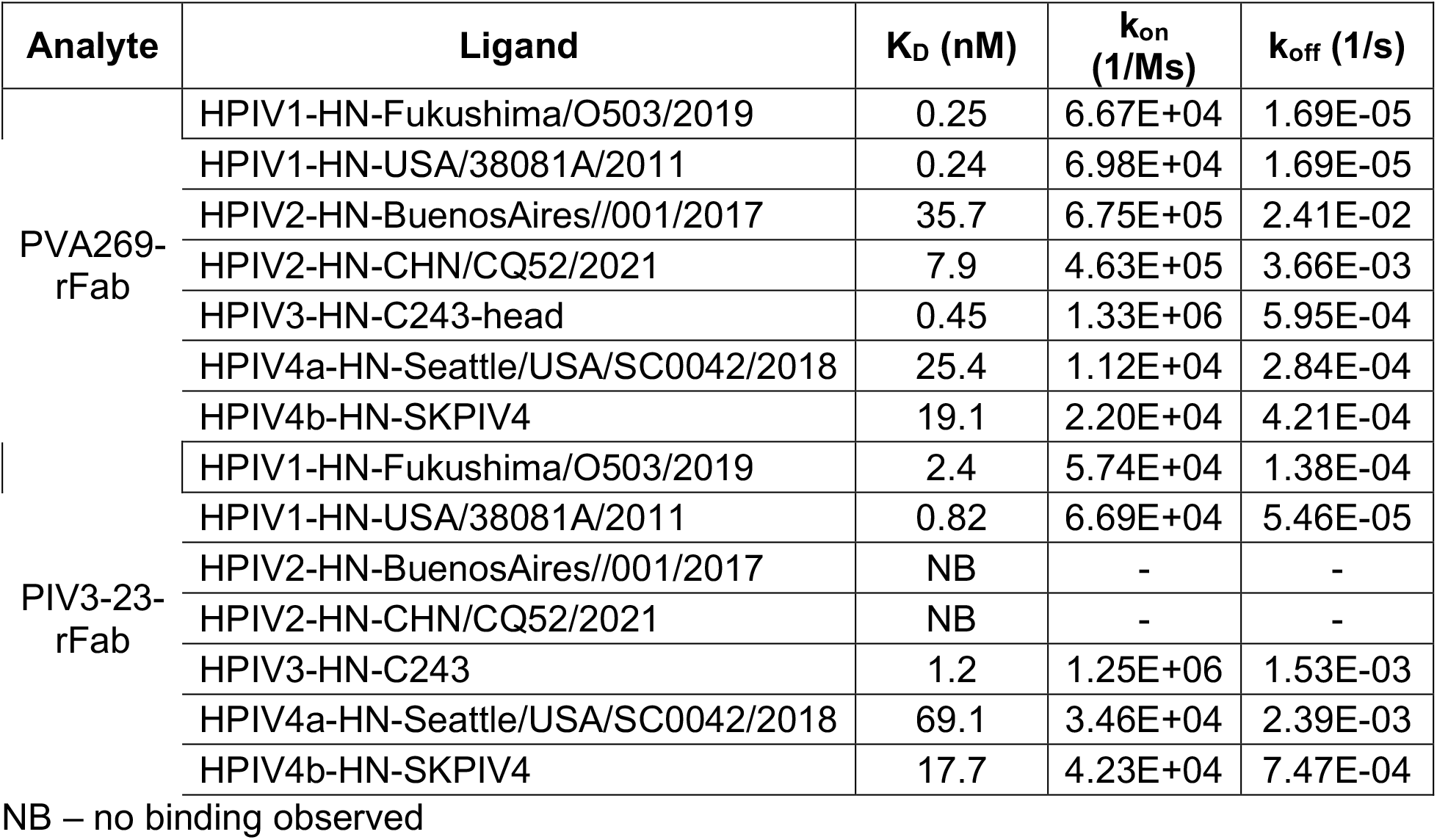
Kinetics parameters of PVA269 or PIV3-23 recombinant Fabs to different HPIV-HN antigens from SPR binding assays.

| Analyte | Ligand | K<sub>D</sub> (nM) | k<sub>on</sub> (1/Ms) | k<sub>off</sub> (1/s) |
| --- | --- | --- | --- | --- |
| PVA269-rFab | HPIV1-HN-Fukushima/O503/2019 | 0.25 | 6.67E+04 | 1.69E-05 |
|  | HPIV1-HN-USA/38081A/2011 | 0.24 | 6.98E+04 | 1.69E-05 |
|  | HPIV2-HN-BuenosAires//001/2017 | 35.7 | 6.75E+05 | 2.41E-02 |
|  | HPIV2-HN-CHN/CQ52/2021 | 7.9 | 4.63E+05 | 3.66E-03 |
|  | HPIV3-HN-C243-head | 0.45 | 1.33E+06 | 5.95E-04 |
|  | HPIV4a-HN-Seattle/USA/SC0042/2018 | 25.4 | 1.12E+04 | 2.84E-04 |
|  | HPIV4b-HN-SKPIV4 | 19.1 | 2.20E+04 | 4.21E-04 |
| PIV3-23-rFab | HPIV1-HN-Fukushima/O503/2019 | 2.4 | 5.74E+04 | 1.38E-04 |
|  | HPIV1-HN-USA/38081A/2011 | 0.82 | 6.69E+04 | 5.46E-05 |
|  | HPIV2-HN-BuenosAires//001/2017 | NB | - | - |
|  | HPIV2-HN-CHN/CQ52/2021 | NB | - | - |
|  | HPIV3-HN-C243 | 1.2 | 1.25E+06 | 1.53E-03 |
|  | HPIV4a-HN-Seattle/USA/SC0042/2018 | 69.1 | 3.46E+04 | 2.39E-03 |
|  | HPIV4b-HN-SKPIV4 | 17.7 | 4.23E+04 | 7.47E-04 |

NB – no binding observed

**Table S2.** CryoEM data collection, processing and model building statistics for the PVA269 Fab-bound HPIV2 HN structure.

|  |  |  |
| --- | --- | --- |
|  | **PVA269-bound**<br>**HPIV2 HN dimer**<br>**PDB 37JG**<br>**EMD 78229** | **PVA269-bound**<br>**HPIV2 HN tetramer**<br>**PDB 37JH**<br>**EMD 78230** |
| **Data collection and processing** |  |  |
| **Magnification** | 105,000 |  |
| **Voltage (kV)** | 300 |  |
| **Electron exposure (e-/Å<sup>2</sup>)** | 30.7 |  |
| **Defocus range (µm)** | -0.6 - -1.6 |  |
| **Pixel size (Å)** | 0.829 |  |
| **Symmetry imposed** | C2 |  |
| **Initial particle images (no.)** | 2,211,257 |  |
| **Final particle images (no.)** | 370,377 | 470,764 |
| **Map resolution (Å)**<br>**FSC threshold** | 2.6<br>0.143 | 2.4<br>0.143 |
| **Refinement** |  |  |
| **Model resolution (Å)**<br>**FSC threshold** | 2.8<br>0.5 | 2.7<br>0.5 |
| **Map sharpening *B* factor (Å<sup>2</sup>)** | -93.7 | -82.9 |
| **Model composition** |  |  |
| Non-hydrogen atoms | 11098 | 20374 |
| Protein residues | 1364 | 2552 |
| Ligands | 46 | 86 |
| Water | 292 | 496 |
| ***B* factors (Å<sup>2</sup>)** |  |  |
| Protein | 31.58 | 102.38 |
| Ligand | 69.45 | 152.66 |
| Water | 21.14 | 88.62 |
| **R.m.s. deviations** |  |  |
| Bond lengths (Å) | 0.003 | 0.004 |
| Bond angles (°) | 0.571 | 0.628 |
| **Validation** |  |  |
| MolProbity score | 1.58 | 1.46 |
| Clashscore | 6.53 | 5.36 |
| Poor rotamers (%) | 0.60 | 0.70 |
| **Ramachandran plot** |  |  |
| Favored (%) | 96.59 | 97.02 |
| Allowed (%) | 3.41 | 2.98 |
| Disallowed (%) | 0 | 0 |

**Table S3.**
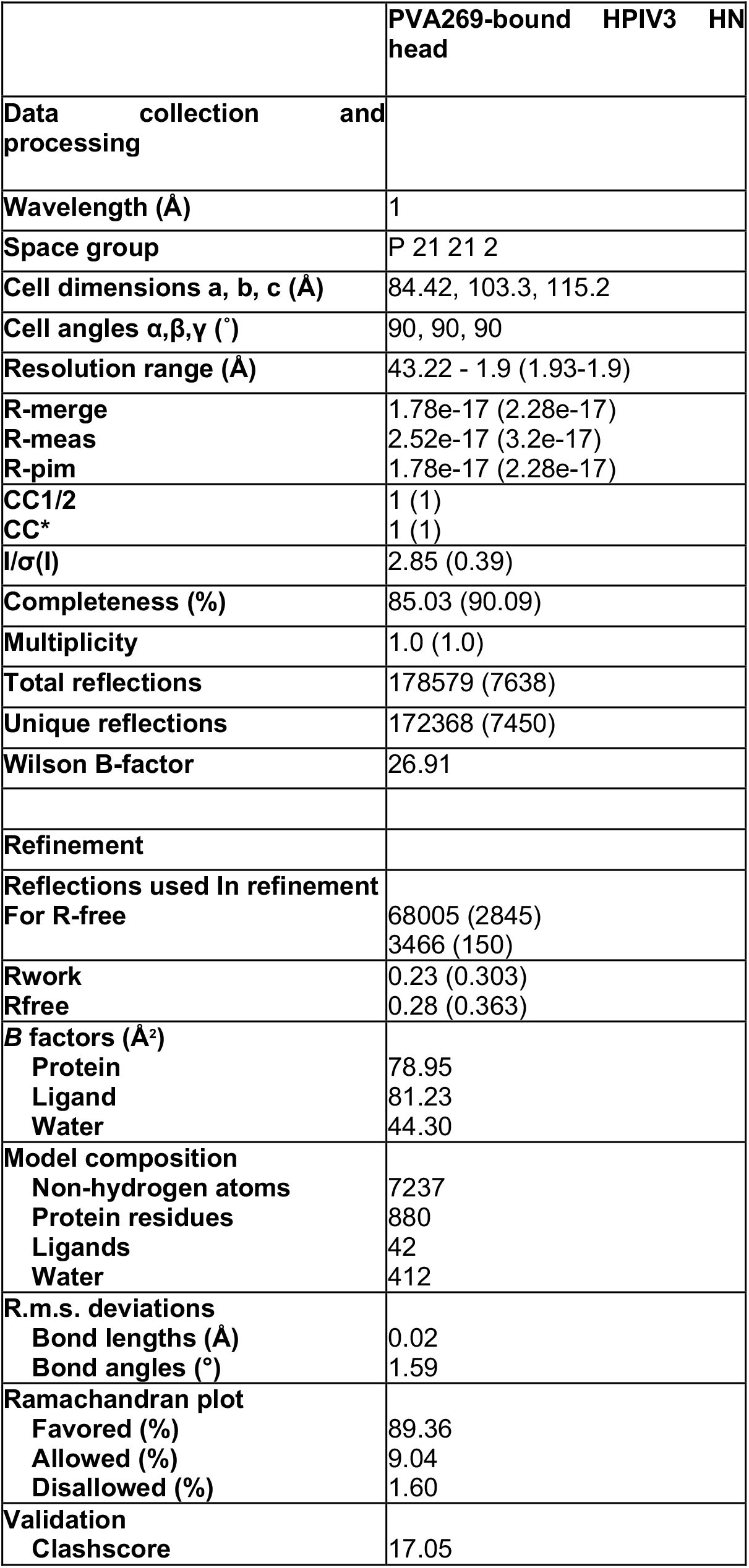
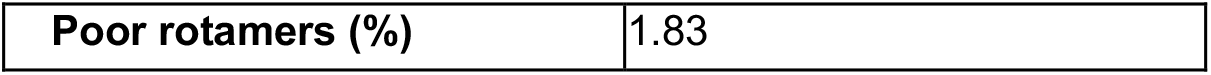
X-ray crystallography data collection, processing and model building statistics for the PVA269 Fab-bound HPIV3 HN head structure.

**Table S4.** BLI K_D_, on rate (Kon), and off rates (K_off)_ of two biological replicates of HPIV2 HN wildtype (WT) or two technical replicates of R354A mutant with PVA269 IgG or Fab. ND = not determined due to very weak binding.

| Ligand | Analyte | K<sub>D</sub> (nM) | k<sub>on</sub> (1/Ms) | k<sub>off</sub> (1/s) |
| --- | --- | --- | --- | --- |
| HPIV2 HN WT | IgG | 0.017 | 5.47E+05 | 9.34E-06 |
|  |  | 0.015 | 7.05E+05 | 1.07E-05 |
|  | Fab | 15.35 | 5.41E+05 | 6.93E-03 |
|  |  | 17.42 | 4.60E+05 | 8.00E-03 |
| HPIV2 HN R354A | IgG | 0.686 | 5.23E+05 | 3.59E-04 |
|  |  | 0.647 | 5.53E+05 | 3.58E-04 |
|  | Fab | ND | ND | ND |

## Acknowledgements

This work was supported in part by the NIH/NIAID under grants 1U19AI181881 and 75N93022C00036 to D.V., an Investigators in the Pathogenesis of Infectious Disease Awards from the Burroughs Wellcome Fund to D.V., and the University of Washington Arnold and Mabel Beckman Cryo-EM Center. D.V. is an investigator of the Howard Hughes Medical Institute, and the Hans Neurath Endowed Chair in Biochemistry at the University of Washington.

## Methods

### Cells and viruses

LLC-MK2 cell line was obtained from the American Type Culture Collection (ATCC). Cells were maintained in MEM, GlutaMAX™ medium supplemented with 10% v/v fetal bovine serum (FBS) and 50 U/ml penicillin/50 µg/ml streptomycin. CHO cells were obtained from ThermoFisher and maintained in ExpiCHO Expression Medium (Thermo Fisher Scientific).

Parainfluenza viruses were purchased from the following vendors:

1. Viratree: HPIV1-GFP – derived from Washington/20993/1964, HPIV2-GFP – derived from Vanderbilt/1994, HPIV3-GFP – derived from JS strain, Sendai virus, PIV5 (W3A strain);
2. BEI: HPIV1/FRA/29221106/2009, HPIV1/FRA/27344044/2007, HPIV2 Greer, HPIV3/Wash/47885/57 (NIH47885), HPIV4a/M25, HPIV4b/CH19503;
3. ATCC: HPIV1/C35, HPIV3/C243, HPIV3/ATCC-2011-5

### Tonsillar sample donors

Tonsils were obtained from individuals undergoing programmed tonsillectomy in local hospitals following approval by the Canton Ticino Ethics Committee (Switzerland). Individuals provide informed consent for the use of biological material for research purposes. Samples were anonymized and no donor-identifying information was made available to the investigators.

### Tonsillar organs processing for cryopreservation

Freshly received tonsils were finely minced with sterile surgical tools and enzymatically digested for 1-2 hours at 37°C with collagenase (2 mg/ml) and DNase I (1 mg/ml) in RPMI 1640 medium supplemented with 1% FBS, 10 mM HEPES, penicillin-streptomycin, kanamycin, amphotericin B. Supernatant was collected, remaining organ fragments mechanically disrupted using a syringe plunger and filtered through 100 µm cell strainers. Cell suspensions from multiple extraction rounds were pooled and centrifuged at 400×g for 7 min at room temperature. The resulting pellet was washed once in supplemented RPMI medium and mononuclear cells were isolated by density-gradient centrifugation over Ficoll-Paque PLUS (400×g, 20 min, room temperature, without brake). Cells were washed, counted, and cryopreserved in FBS containing 10% DMSO before storage in liquid nitrogen.

### Construct design

To generate the HPIV2 HN ectodomain construct, residues 47-571 of human parainfluenza virus 2 hemagglutinin-neuraminidase (Genbank: NP_598405.1) were cloned into a pCMV vector with an N-terminal 8xHis tag and a flexible GGSG linker. The native signal peptide was replaced with the Mu-phosphatase signal peptide. The R354A mutant construct was generated via site-directed mutagenesis. All plasmids were synthesized by Genscript.

The other HN proteins (head or ectodomain) were produced by VIVA Biotech.

### Purification of recombinant HPIV2 HN

100 mL of Expi293 cells were transfected with HPIV2 HN wildtype or R354A mutant construct following the Expi293 manual. Five days post transfection, cells were harvested by centrifugation at 3500 rpm for 30 minutes, and the resulting lysate was clarified by vacuum filtration through a 0.22 µm membrane. The clarified lysate was applied to 3 mL of Ni Sepharose Excel resin slurry pre-equilibrated in base buffer (25 mM sodium phosphate, 300 mM NaCl, pH 8.0) and incubated with gentle agitation at 155 rpm and 20°C to allow batch binding. The resin was then washed with 10 column volumes of wash buffer (base buffer supplemented with 20 mM imidazole), and bound protein was eluted with 20 mL of elution buffer (base buffer supplemented with 500 mM imidazole).

The eluate was subjected to size-exclusion chromatography at room temperature using a Superdex 200 Increase 10/300 GL column equilibrated in SEC buffer (50 mM Tris-HCl, 150 mM NaCl, pH 8.0). Peak fractions were assessed by SDS-PAGE, and those containing the target protein were pooled for downstream use.

### Antigen-specific memory B cell repertoire analysis (AMBRA)

Total mononucleated tonsillar cells were seeded into round bottomed 96-well plates (20,000 cells/well in 200 ul) in RPMI-1640 supplemented with 10% FBS, 1 mM sodium pyruvate, 1% MEM NEAA solution, 1% Glutamax, 50 mm b-mercaptoethanol, 50 IU penicillin/50 ug/ml streptomycin, 1% Kanamycin, 2.5 mg/ml R848 and 1000 IU/ml human recombinant IL-2. Cells were incubated at 37°C 5% CO_2_. After 10 days, culture supernatant was collected and stored at -20°C until further processing.

### Enzyme-Linked Immunosorbent Assay (ELISA) for tonsillar donor selection

HPIV1-4 HN proteins were prepared at 1 mg/ml in PBS (pH 7.2) and absorbed onto 384- well plates (5 ml/well) overnight at 4°C. Plates were washed with PBS 0.05% Tween 20 (PBS-T) with a microplate washer (Agilent Biotek) and blocked with PBS 1% BSA (20 ml/well) for at least 60 min at room temperature. Blocking solution was aspirated and 5 ml/well AMBRA supernatant dispensed using an automatized robotic liquid handler. After 60 min incubation at room temperature, plates were washed twice with PBS-T goat anti-human IgG secondary antibody added and incubated for 60 minutes at room temperature. Plates were then washed again with PBS-T and 4-NitroPhenyl phosphate (pNPP) substrate added. After 30 min incubation, absorbance at 405 nm was measured by a plate reader (Agilent Biotek) and data exported and analyzed in Excel.

### Antibody isolation and recombinant production

Starting for cryopreserved mononucleated cells, B cells were enriched via PE-Cy7-anti CD19 staining 20 min at 4°C, followed by a washing step in PBS 2% FBS 2mM EDTA and anti-PE beads staining for 20 min at 4°C for positive selection using LS columns (Miltenyi). Enriched B cells were stained with PE labelled anti-IgM, anti-IgD, anti-CD14 and anti-IgA and biotinylated HPIV2-HN previously conjugated with AlexaFluor 647-Streptavidin. After 20 min incubation at 4°C, cells were washed and antigen specific memory B cells were isolated with fluorescence-activated cell sorting (FACS). Selected cells were seeded clonally into 384-well plates on a monolayer of mesenchymal stem cells in 384-well microtiter plates in the presence of stimulation medium (IMDM 10% FBS, 2.5 μg/ml CpG 2006, 500 IU/ml IL-2, 5 ng/ml IL-6 (BD Pharmingen), 50 ng/ml IL-10 (ImmunoTools) and 10 ng/ml IL-21 (ImmunoTools). After 7 days of incubation at 37°C 5% CO_2_, B cells supernatants were screening for binding capacity to HPIV3-HN by ELISA and for neutralization activity against HPIV2 by laser scanning cytometry (Mirrorball® - TPP LABTECH).

Heavy-and light-chain variable region genes were amplified from B-cell cDNA by nested PCR using Q5 High-Fidelity DNA Polymerase (New England Biolabs). Purified amplicons were cloned into the human IgG1m17 or IgG1m3 and human pkappa expression vectors. Vector-insert assemblies were generated using In-Fusion Snap Assembly Master Mix (Takara Bio) according to the manufacturer’s instructions and transformed into Zymo 10B competent *Escherichia coli* cells. Transformants were selected on ampicillin-containing agar plates and screened by Sanger sequencing using a vector-specific forward primer. Codon-optimized plasmids were synthetized for expression in Chinese hamster ovary (CHO) cells as IgG1m17 or Fab as follows: VH-and VL-coding plasmids were co-transfected into CHO cells using the ExpiFectamine CHO Transfection Kit (Thermo Fisher Scientific). Cultures were supplemented 24 h post-transfection and maintained for 8 days before harvest. Clarified supernatants were collected by 0.22-µm filtration and subjected to antibody purification. Monoclonal antibodies were purified by affinity chromatography on an ÄKTA Xpress FPLC system (Cytiva) operated with UNICORN software v5.11 (Build 407). Full-length human, whereas Fab fragments were purified using CaptureSelect CH1-XL MiniChrom columns (Thermo Fisher Scientific). Phosphate-buffered saline (PBS) was used as the mobile phase. Purified proteins were buffer-exchanged into the appropriate formulation buffer using HiTrap Fast Desalting columns (Cytiva), sterile-filtered through 0.22-µm membranes, and stored at 4°C.

### PIV microneutralization assay by laser scanning cytometry

This protocol was applied for the screening of stimulated B cell culture supernatant and the characterization of purified monoclonal antibodies against HPIV1-GFP (Washington/20993/1964), HPIV2-GFP (Vanderbilt/1994), HPIV3-GFP (JS strain), Sendai virus and PIV5 (W3A strain).

Virus stocks were propagated on LLC-MK2 cells as follows: in a T75 flask format, cell were put in culture with an input virus at 0.01 MOI in infection medium (MEM, GlutaMAX™ 3% v/v FBS and 50 U/ml penicillin/50 µg/ml streptomycin). To allow spreading, TPCK-treated trypsin prepared in infection medium at 55 mg/ml (for HPIV1 and HPIV2) or 35 mg/ml (for Sendai virus) was added 24 hours of viral/cell culture. After 3 to 5 days, cell culture supernatant was collected, and cells scraped and lysed by snap-freezing. Supernatant were pooled and cleared by centrifugation at 1500xg for 5 min at 4°C. Sucrose 5% and HEPES 50 mM were added, solution aliquoted, snap frozen and stored at -80°C until use. For virus stock titration, LLC-MK2 cells were infected in a 384-well plate format with serial virus dilution in infection medium and incubated at 37°C 5% CO_2_ for 72 hours. After the incubation time, cells were stained with 3 nM Draq5 in infection medium for 4 hours at 37°C and plates read at the Mirrorball scanner cytometer. Data were exported in Excel as percentage of infected cells (GFP positive cells in the Draq5 positive population) and virus titer calculated according to the Spearman-Karber formula.

For the neutralization assays, 10 ml of stimulated B cell culture supernatant or recombinant antibodies titrations prepared in infection medium were dispensed into black, clear bottom 384-well plates using a robotic liquid handler or manually, respectively. Input viruses were prepared in infection medium (containing only 1% FBS from primary screening of B cell supernatants) in order to infect cells with 250 TCID50/well. After 45 min of antibody/virus incubation at 37°C 5% CO_2_, 1200 LLC-MK2 cells/well were added in infection medium and plates placed back in the incubator for the following 72 hours. TPCK-treated trypsin was added at the conditions described above, if required. At the time of read out, cells’ nuclei were stained with 3 nM Draq5 (ThermoFisher) in infection medium for at least 4 hours at 37°C 5% CO_2_. Plates were read using a Mirrorball® scanner cytometer using Draq5 signal to identify cells. Data were exported in Excel as percentage of infected cells (GFP positive cells in the Draq5 positive population). For monoclonal antibodies analysis data were plotted in GraphPad Prism Software (v11.0.2) The IC_50_ values were calculated using a non-linear regression model (variable slope model, four parameters) fitting model.

### HPIV neutralization assay (96-well plate format)

This protocol was used to characterize the neutralization potency of purified monoclonal antibodies against the virus panel reported in Fig.1D.

Virus stocks were propagated on LLC-MK2 cells as follows: 2.5 million cells were seeded in a T75 flask in EMEM GlutaMAX™ 10% v/v FBS 1% NEAA 50 U/ml penicillin/50 µg/ml streptomycin and incubated overnight at 37°C. After a washing step, cells were inoculated with HPIVs prepared at 0.001 or 0.1 MOI in infection medium (EMEM, GlutaMAX™ 1% v/v or 2.4% FBS and 50 U/ml penicillin/50 µg/mL streptomycin) for 1 hour at 37°C. To facilitate spreading of HPIV1 and HPIV4 strains, infection medium was supplemented with 25 μg/ml TPCK-treated trypsin. After 3 to 10 days, cell culture supernatant was collected, cells scraped and lysed by snap-freezing. Supernatants were pooled and cleared by sequential centrifugation steps at 500xg and at 2000xg for 5 min at 4°C each. For virus stock titration, LLC-MK2 cells were seeded into 96-well flat bottom tissue culture plates at 20,000 cells/well and incubated overnight at 37°C. Twenty-four hours later, complete medium was aspirated and cells were infected with two-fold serial dilutions of stock viruses for 30 minutes at 37°C. After virus adsorption, cells were washed and overlaid with100 μl/well of infection media 2.4% colloidal cellulose. After 48 hours incubation at 37°C 5% CO_2_, cells were fixed with 4% paraformaldehyde, permeabilized with permeabilization buffer (0.5% Triton-X 100 in PBS) and stained with anti-HN antibodies specific for each HPIV subtype in wash/perm buffer followed by goat anti-mouse IgG Alexa Fluor Plus 647 along with Hoechst. The plates were washed and imaged on the Ensight Multimode Plate Reader (PerkinElmer) and virus titers were calculated according to the Spearman-Karber formula.

For the neutralization assay, LLC-MK2 cells were seeded into a flat-bottom 96-well tissue culture plates at 20,000 cells /well and cultured overnight at 37°C in growth medium. A 1:1 mix of purified mAbs serial dilutions and input viruses (100 FFU/well) was prepared in infection media (containing only 1% or 2.4% FBS for HPIV2/HPIV3 strains and HPIV1/HPIV4 strains, respectively) and incubated at 37°C 5% CO_2_, for 1 hour.

Antibody/virus mixtures were added to the cells and incubated at 37°C for 48 hours. Cells were then fixed, stained and imaged using the same procedure described above for the virus titration assay. The data was normalized by calculating the percent neutralization relative to the untreated control after correction for the background signal. Neutralization curves were plotted in GraphPad Prism Software (v11.0.2) and IC50 values calculated using a non-linear regression model (variable slope model, four parameters) fitting model.

### Antibody affinity determination by Surface Plasmon Resonance (SPR)

Measurements were performed using a Biacore T200 instrument. Recombinant HPIV-HN antigens containing Avi-Tags were biotinylated with BirA biotin-protein ligase according to manufacturer’s protocol (Avidity LLC). The Biotin CAPture kit (Cytiva) was used to capture biotinylated HN proteins. Running buffer was HBS-EP+ pH 7.4 (Cytiva) and measurements were performed at 25°C. Single cycle kinetics experiments were performed with a 5-fold dilution series of PVA269 or PIV3-23 recombinant Fabs: 1.17, 4.69, 18.75, 75.0, 300 nM. Data were double reference-subtracted and fit to a binding model using Biacore Evaluation software. The 1:1 binding model was used to estimate the kinetic parameters.

### MUNANA assay

mAbs were serially diluted in assay buffer (1% BSA in PBS +Ca/Mg) and mixed in 96-well black plates (Greiner) with a final concentration of recombinant HN glycoprotein (VIVA Biotech Limited) leading to 2x10^6^ RFU signal as determined by a previous titration step and summarized here below:

-HPIV1_USA/38081A/2018-HN head: 41.7 ng/ml

- HPIV1_Fukushima/O503/2019-HN head: 105.7 ng/ml

- HPIV2_CHN/CQ52/2021-HN ecto: 3.9 ng/ml

- HPIV3_C243-HN head: 1451.7 ng/ml

- HPIV4a_Seattle/USA/SC0042/2018-HN ecto: 401.6 ng/ml

After an incubation of 30 minutes at 37°C, MUNANA substrate (Sigma Aldrich) prepared in MUNANA buffer (10 mM CaCl2, 100 mM sodium acetate pH 4.5), was added at a final concentration of 200 μM and plates were incubated for 2 h at 37 °C. Enzymatic reaction was stopped with MUNANA stop solution (200 mM sodium carbonate pH 9.5) and fluorescence measured using Cytation5 reader (Agilent Biotek) (excitation at 365 nm and emission at 450 nm). Data were analyzed in GraphPad Prism (v11.0.2) and mAbs IC50 values calculated using a non-linear regression model (variable slope model, four parameters) fitting model.

### Hemagglutination inhibition (HI) assay

HPIV1-3 hemagglutination unit (HAU) titer was determined by incubating 50 μl of 0.75% guinea pig red blood cell suspension with 50 μl of two-fold serially diluted HPIV virus stocks (HPIV1, HPIV2, and HPIV3) in V-bottom 96-well plates for 1 hour at room temperature. Viral dilution leading to 4 HA units was calculated for each. For HI assay, 25 μl of mAbs two-fold serial dilutions (prepared in ice-cold PBS) were incubated with 25 μl containing 4 HAU of the different HPIV virus strains for 1 hour at room temperature in V-bottom 96-well plates. Fifty μl of 0.75% guinea pig red blood cells diluted in ice-cold PBS was added to the virus:mAb mixture and incubated for 1 hour at room temperature. The IC100 values were calculated based on the lowest concentration of mAb that inhibited the hemagglutination of the guinea pig red blood cells.

### Biolayer Interferometry

BLI assays were performed on an Octet Red instrument (Sartorius) at 30°C and 1,000 rpm. Ni-NTA biosensors were pre-hydrated for 10 minutes in 1X kinetics buffer made by diluting 10X kinetics buffer (Sartorius) in PBS prior to use. For KD determination of HPIV2 HN WT/R354A and PVA269 IgG, His-tagged HN was loaded onto Ni-NTA biosensors (Sartorius, Cat# 18-5101) at 25 µg/mL in 1X KB until a 1 nm shift was reached and then dipped into 1X KB for 60 sec before 600 sec association with 6.67 to 0.247 nM of IgG, serially diluted threefold. Dissociation followed in 1X KB for 1,800 sec for the WT HN and IgG, and 1,200 sec for all others. The same setup was used for HPIV2 HN WT/R354A and PVA269 Fab, except the concentration range of Fab used was 160 to 1.98 nM, serially diluted threefold. Sensorgrams were adjusted by subtraction of a reference loaded biosensor dipped into 1X KB for the association and dissociation phases. Association, dissociation, and KD values were calculated with ForteBio data analysis software with a 1:1 binding model and global fit of curves.

### Cryo-EM Sample Preparation, Data Collection, and Data Processing

HPIV2 HN and PVA269 complex was prepared by mixing a 1:5 molar ratio of HN:PVA269 Fab followed by a 1h incubation at 4°C. 3 uL of 2.24 µM complex was applied onto fresh glow discharged R 2/2 UltrAuFoil grids prior to plunge freezing using a Vitrobot MarkIV (ThermoFisher Scientific) with a blot force of 0, 2.5 and 4.5 sec sec blot time, and 40 sec waiting time at 100% humidity. The data was acquired using an FEI Titan Krios transmission electron microscope operated at 300 kV and equipped with a Gatan K3 direct detector and Gatan Quantum GIF energy filter, operated in zero-loss mode with a slit of 20 eV. Automated data collection was carried out using SerialEM at a nominal magnification of 105,000x with a pixel size of 0.829 Å. A total of 35,464 micrographs were collected from two sessions with a defocus range between -0.6 and -1.6 μm and stage tilt angle of 0°. Movie frame alignment, estimation of the microscope contrast-transfer function parameters, particle picking, and extraction were carried out using cryoSPARC Live^38^ and all downstream processing steps were done in cryoSPARC. Particles were extracted with a box size of 512 pixels and downsampled by a factor of 2. Two rounds of reference-free 2D classification were performed to select well-defined particle images, and here we noticed two populations of HPIV2 HN, consisting of dimer and tetramer heads. Initial model generation for dimer and tetramer populations were performed using ab-initio reconstruction and the resulting maps were used for homogeneous and subsequent heterogeneous refinement. Re-extracted particles (272K for tetramer and 540K for dimer) were used for subsequent non-uniform refinement with per-particle defocus and global CTF refinement. Three rounds of topaz training and extraction followed to improve particle picks, which led to 479K particles for the tetramer and 376K for the dimer after downstream processing. Reference-based motion correction and final non-uniform refinement with per-particle defocus and global CTF refinement was performed to obtain final reconstructions of the dimer and tetramer HN bound to PVA269 at 2.6Å and 2.4Å respectively (EMD: 78229 and EMD: 78230). Reported resolutions are based on the gold-standard Fourier shell correlation (FSC) of 0.143 criterion. See also Fig S4.

### Model building

ModelAngelo^39^ was used for initial model building with sequences of the PVA269 IgG and HPIV2 HN ectodomain. The structure was then manually rebuilt using Coot^40^ and refined using Phenix^41^. Validation used Molprobity^42^, Phenix, and Privateer^43^. Structures are deposited in the PDB with accession codes 37JG and 37JH for the dimer and tetramer complex structures respectively.

### Conservation of PVA269 epitope

A logo plot for amino acid conservation of PVA269 epitope, created using Logomaker^44^, was based on available HN sequences from HPIV2 (n = 210) and HPIV3 (n = 2140). These publicly available sequences were downloaded from NCBI using an upper limit date cutoff of 2023-08-23. The earliest collected sequence was used as the reference for each subtype. Sequences were aligned using MAFFT version 7^45,46^ with automatically determined setting (–auto). Sequences with > 10 consecutive nucleotide deletions were removed. Frequency in the logo plot is based on sequences where an amino acid is found to be aligned in each of the reported epitope-relevant positions.

### Animals and HPIV3 challenge

6-8 weeks old female *Sigmodon hispidus* were maintained and handled under veterinary supervision following the National Institutes of Health guidelines and Sigmovir’s Institutional Animal Care and Use Committee-approved animal study protocol. Cotton rats were housed in clear polycarbonate cages and provided standard rodent chow (Harlan #7004) and tap water *ad lib.* Animals in six groups (N=8) were inoculated with mAb intramuscularly 1 day prior to intranasal infection with 100 μl of 10^5^ PFU HPIV3 per animal. 4 days post infection, nasal turbinates and lungs were collected for viral titration. Sera was collected on day 0 (before infection) to measure PVA269 concentration.

### ELISA for IgG quantification in sera

Human IgG concentrations were determined using an electrochemiluminescence immunoassay. Standards and quality control samples (QCs) were prepared in pooled Sprague Dawley rat serum (BioIVT), aliquoted, and stored at −80 °C until use. Reagents and samples were equilibrated to room temperature for ≥30 min before use.

Samples, standards, and QCs were centrifuged at 1,000 × g for 1 min before analysis. Multi-array 96-well plates (Meso Scale Discovery) were coated with heavy-chain-specific polyclonal goat anti-human IgG (SouthernBiotech; 2 μg/ml) for 120 min at room temperature with shaking (650 rpm), washed with PBS-T, and blocked with 5% bovine serum albumin (BSA) in PBS for 1 h.

Serum samples were diluted in pooled Sprague Dawley rat serum as appropriate for each sampling time point. Samples, standards, and QCs were subsequently diluted 1:30 in 1% BSA in PBS as the minimum required dilution (MRD).

MRD samples, standards, and QCs were added to the plates (50 μl/well) and incubated for 1 h at room temperature. After washing, ruthenylated F(ab′)₂-specific goat anti-human IgG (Jackson ImmunoResearch; 0.125 μg/ml) was added for detection of PVA269-rIgG1m3 and incubated for 1 h. Plates were washed, MSD Read Buffer was added (150 μl/well), and signals were acquired using a MESO QuickPlex SQ 120.

### Plaque assay

Lung and nose homogenates were clarified by centrifugation and diluted in EMEM. Confluent HEp-2 monolayers were infected in duplicates with diluted homogenates in 24 well plates. After 1 hour incubation at 37°C in a 5% CO_2_ incubator, the wells were overlaid with 0.75% Methylcellulose medium. After 4 days of incubation, the overlay was removed and the cells were fixed with 0.1% crystal violet stain for one hour and then rinsed and air dried. Plaques were counted and virus titer was expressed as plaque forming units per gram of tissue. Viral titers are calculated as geometric mean ± standard error for all animals in a group at a given time. The limits of detection in the plaque assays are 2.3 log10 PFU/g for the lung, and 2.0 log10 PFU/g for the nose viral titers.

### Statistical analyses

All statistical tests were performed as described in the indicated figure legends using Prism v10.0. The number of independent experiments performed is indicated in the relevant figure legends.

## References

1. Hall, C. B. Respiratory syncytial virus and parainfluenza virus. N. Engl. J. Med. 344, 1917–1928 (2001).

2. Henrickson, K. J. Parainfluenza viruses. Clin. Microbiol. Rev. 16, 242–264 (2003).

3. CDC. Clinical overview of human Parainfluenza viruses. Parainfluenza https://www.cdc.gov/parainfluenza/hcp/clinical-overview/index.html (2026).

4. Fry, A. M. et al. Seasonal trends of human parainfluenza viral infections: United States, 1990-2004. Clin. Infect. Dis. 43, 1016–1022 (2006).

5. Ustun, C. et al. Human parainfluenza virus infection after hematopoietic stem cell transplantation: risk factors, management, mortality, and changes over time. Biol. Blood Marrow Transplant. 18, 1580–1588 (2012).

6. Ogimi, C. et al. Novel factors to predict respiratory viral disease progression in allogeneic hematopoietic cell transplant recipients. Bone Marrow Transplant. 57, 649–657 (2022).

7. Tabatabai, J. et al. Parainfluenza virus infections in patients with hematological malignancies or stem cell transplantation: Analysis of clinical characteristics, nosocomial transmission and viral shedding. PLoS One 17, e0271756 (2022).

8. Pawełczyk, M. & Kowalski, M. L. The role of human Parainfluenza virus infections in the immunopathology of the respiratory tract. Curr. Allergy Asthma Rep. 17, 16 (2017).

9. Lawrence, M. C. et al. Structure of the haemagglutinin-neuraminidase from human parainfluenza virus type III. J. Mol. Biol. 335, 1343–1357 (2004).

10. Yuan, P. et al. Structural studies of the parainfluenza virus 5 hemagglutinin-neuraminidase tetramer in complex with its receptor, sialyllactose. Structure 13, 803– 815 (2005).

11. Porotto, M. et al. Regulation of paramyxovirus fusion activation: the hemagglutinin-neuraminidase protein stabilizes the fusion protein in a pretriggered state. J. Virol. 86, 12838–12848 (2012).

12. Marcink, T. C. et al. How a paramyxovirus fusion/entry complex adapts to escape a neutralizing antibody. Nat. Commun. 15, 8831 (2024).

13. Moscona, A. Entry of parainfluenza virus into cells as a target for interrupting childhood respiratory disease. J. Clin. Invest. 115, 1688–1698 (2005).

14. Suryadevara, N. et al. Functional and structural basis of human parainfluenza virus type 3 neutralization with human monoclonal antibodies. Nat. Microbiol. 9, 2128– 2143 (2024).

15. Domachowske, J. et al. Safety of nirsevimab for RSV in infants with heart or lung disease or prematurity. N. Engl. J. Med. 386, 892–894 (2022).

16. Stewart-Jones, G. B. E. et al. Structure-based design of a quadrivalent fusion glycoprotein vaccine for human parainfluenza virus types 1-4. Proc. Natl. Acad. Sci. U. S. A. 115, 12265–12270 (2018).

17. McCaffrey, K. D., Esfahani, B. G., Elbehairy, M. A., McCormick, A. L. & Mousa, J. J. Molecular basis for protection and cross-protection by human antibodies targeting the parainfluenza virus hemagglutinin-neuraminidase protein. J. Virol. e0050226 (2026).

18. Cabán, M. et al. Cross-protective antibodies against common endemic respiratory viruses. Nat. Commun. 14, 798 (2023).

19. Miller, R. J. et al. The structural basis of protective and nonprotective human monoclonal antibodies targeting the parainfluenza virus type 3 hemagglutinin-neuraminidase. Nat. Commun. 15, 10825 (2024).

20. Foglierini, M., Pappas, L., Lanzavecchia, A., Corti, D. & Perez, L. AncesTree: An interactive immunoglobulin lineage tree visualizer. PLoS Comput. Biol. 16, e1007731 (2020).

21. Welch, B. D. et al. Structure of the parainfluenza virus 5 (PIV5) hemagglutinin-neuraminidase (HN) ectodomain. PLoS Pathog. 9, e1003534 (2013).

22. Wang, Z. et al. Architecture and antigenicity of the Nipah virus attachment glycoprotein. Science 375, 1373–1378 (2022).

23. Wang, Z. et al. Structure and design of Langya virus glycoprotein antigens. Proc. Natl. Acad. Sci. U. S. A. 121, e2314990121 (2024).

24. Yuan, P. et al. Structure of the Newcastle disease virus hemagglutinin-neuraminidase (HN) ectodomain reveals a four-helix bundle stalk. Proc. Natl. Acad. Sci. U. S. A. 108, 14920–14925 (2011).

25. Kalbermatter, D. et al. Structure and supramolecular organization of the canine distemper virus attachment glycoprotein. Proc. Natl. Acad. Sci. U. S. A. 120, e2208866120 (2023).

26. Punjani, A., Rubinstein, J. L., Fleet, D. J. & Brubaker, M. A. cryoSPARC: algorithms for rapid unsupervised cryo-EM structure determination. Nat. Methods 14, 290–296 (2017).

27. Hunt, A. C. et al. Multivalent designed proteins neutralize SARS-CoV-2 variants of concern and confer protection against infection in mice. Sci. Transl. Med. 14, eabn1252 (2022).

28. Lee, J. et al. The computationally designed TRI2-2 miniprotein inhibitor protects against multiple SARS-CoV-2 Omicron variants. *Commun*. Biol. 9, 224 (2026).

29. Ragotte, R. J. et al. Designed miniproteins potently inhibit and protect against MERS-CoV. Cell Rep. 44, 115760 (2025).

30. Abu-Shmais, A. A. et al. Potent HPIV3-neutralizing IGHV5-51 antibodies identified from multiple individuals show L chain and CDRH3 promiscuity. J. Immunol. 212, 1450–1456 (2024).

31. Boonyaratanakornkit, J. et al. Protective antibodies against human parainfluenza virus type 3 infection. MAbs 13, 1912884 (2021).

32. Marcink, T. C. et al. Subnanometer structure of an enveloped virus fusion complex on viral surface reveals new entry mechanisms. Sci. Adv. 9, eade2727 (2023).

33. Acciani, M. et al. Human neutralizing antibodies targeting the measles virus hemagglutinin and fusion surface proteins. Cell Host Microbe 34, 1067–1081.e12 (2026).

34. Momont, C. et al. A pan-influenza antibody inhibiting neuraminidase via receptor mimicry. Nature 618, 590–597 (2023).

35. Stadlbauer, D. et al. Broadly protective human antibodies that target the active site of influenza virus neuraminidase. Science 366, 499–504 (2019).

36. Johnson, S. et al. Development of a humanized monoclonal antibody (MEDI-493) with potent in vitro and in vivo activity against respiratory syncytial virus. J. Infect. Dis. 176, 1215–1224 (1997).

37. Zhu, Q. et al. A highly potent extended half-life antibody as a potential RSV vaccine surrogate for all infants. Sci. Transl. Med. 9, eaaj1928 (2017).

38. Punjani, A. Algorithmic advances in single particle cryo-EM data processing using CryoSPARC. Microsc. Microanal. 26, 2322–2323 (2020).

39. Jamali, K. et al. Automated model building and protein identification in cryo-EM maps. Nature 628, 450–457 (2024).

40. Emsley, P., Lohkamp, B., Scott, W. G. & Cowtan, K. Features and development of coot. Acta Crystallogr. D Biol. Crystallogr. 66, 486–501 (2010).

41. Liebschner, D. et al. Macromolecular structure determination using X-rays, neutrons and electrons: recent developments in Phenix. Acta Crystallogr. D Struct. Biol. 75, 861–877 (2019).

42. Williams, C. J. et al. MolProbity: More and better reference data for improved all-atom structure validation. Protein Sci. 27, 293–315 (2018).

43. Agirre, J. et al. Privateer: software for the conformational validation of carbohydrate structures. Nat. Struct. Mol. Biol. 22, 833–834 (2015).

44. Tareen, A. & Kinney, J. B. Logomaker: beautiful sequence logos in Python. Bioinformatics 36, 2272–2274 (2020).

45. Katoh, K., Misawa, K., Kuma, K.-I. & Miyata, T. MAFFT: a novel method for rapid multiple sequence alignment based on fast Fourier transform. Nucleic Acids Res. 30, 3059–3066 (2002).

46. Katoh, K. & Standley, D. M. MAFFT multiple sequence alignment software version 7: improvements in performance and usability. Mol. Biol. Evol. 30, 772–780 (2013).

